# Dual schema of allergens reveals molecular and evolutionary signatures

**DOI:** 10.64898/2025.12.12.694064

**Authors:** Chunlai Tam

## Abstract

Allergies affect billions of people worldwide, posing a substantial challenge to global health because of its elusive molecular determinants. Leveraging high-quality protein structures, we revisit why allergenic proteins trigger allergies, whereas some of their structural analogues do not. We categorize allergenic protein folds into similar (SAP) and dissimilar (DAP) folds on the basis of their similarity to human proteins, establishing a dual schema that reveals distinct molecular and evolutionary patterns. Here, we show that compared with structurally similar, nonallergenic protein (SNAP) folds, SAP folds are ubiquitous across kingdoms and rich in B- and T-cell epitopes. DAP folds map to a unique sequence–structure–function space and are evolutionarily underrepresented and taxonomically restricted. The persistent divergence of DAP folds from animal proteins suggests their intermittent immune encounters. In contrast, ubiquitous SAP folds require continuous immune sorting from their SNAP analogues. This dichotomy provides a new lens for studying the evolution of host immune responses in allergic reactions. Our classifier, AllerX, is trained on these patterns and predicts unseen APs with state-of-the-art accuracy. This work, which links the structural, immunological, and evolutionary aspects of allergenic proteins, establishes a novel framework for enhancing diagnostics and food safety.

**Significance:** Allergies, a global health challenge, cause life-threatening immune reactions. Our theory links protein foreignness to immunogenicity, but its structural definition based on similarity to human proteins remains undefined. Utilizing massive protein structure resources, we grouped allergenic protein folds into similar (SAP) or dissimilar (DAP) to human proteins. We reveal strong contrasts in molecular and evolutionary signatures: DAP folds are taxonomically restricted but have biased, source-specific distribution among kingdoms. SAP folds show prominent, orientation-specific B-cell epitopes compared to their non-allergenic structural analogs. We also show fold-independent T-cell epitope linkage between SAP and DAP folds as a converged allergenicity signal. This dichotomy enhances cross-reactivity tracing in food safety incidents and establishes a foundation for studying host-factor evolution of protein allergy.

## Introduction

Allergies are a global health burden that affect billions of people and are costly to public health budgets(*1–3*). Allergic diseases can be miserable and even fatal, with relentless symptoms such as sneezing and itching disrupting daily life. The constant fear of anaphylaxis, a life-threatening reaction, can turn every exposure into a potential emergency(*4*).

Allergenic proteins (APs) are a major subject in allergy research(*5*) among various biomolecules(*6–8*) and small molecules(*9*) that trigger allergic reactions. A key prerequisite for allergenicity is the immune system’s recognition of a protein as foreign. Allergens that harbour unique structures often meet this prerequisite, such as the cockroach allergen Bla g 1(*10*). Interestingly, many AP folds, such as EF hand and profilin, which are also present in the human proteome, are evolutionarily ubiquitous. While the molecular determinants of allergenicity for these folds were previously unclear(*11*, *12*), subsequent studies have revealed sequential, structural, and functional differences from their nonallergenic counterparts(*13–22*). However, these studies either relied on a small dataset or focused on a particular fold class or taxonomic lineage of APs, lacking a high-level overview that would lead to a convergent model for AP allergenicity.

More recently, advances in protein structure prediction models(*23*) and the expansion of protein structure databases(*24*, *25*), as well as the development of fast structural search(*26*) and domain segmentation algorithms(*27*, *28*), have unlocked vast opportunities for the investigation of APs at the protein domain level. We argue that studying APs at the domain level is crucial because categorizing their domains into structurally similar (SAP) versus dissimilar (DAP) folds with respect to the human proteome is a critical step in quantifying their “foreignness”, the key metric that lies at the heart of our allergenicity theory. Not dividing into separate domains challenges the categorization of multidomain proteins, which are common. Moreover, as self-standing units of protein evolution, the protein domains identified in APs will aid our understanding of their evolutionary pathways. Utilizing the available resources, we aimed to elucidate the evolutionary history and molecular hallmarks that distinguish SAP folds from DAP folds, as well as from SNAP folds, which are the nonallergenic structural analogues of SAP folds.

We investigated the abundance of AP folds based on structural similarity in the proteomes of animal models (Fig. 1a), including *C. elegans* (CaeEl), *D. melanogaster* (DroMe), *D. rerio* (DanRe), *M. musculus* (MusMu), *P. paniscus* (PanPa), and humans, and discovered a strong bimodal distribution that offered clear segregation into structurally similar and dissimilar folds for each proteome (Fig. S1a). According to this bimodality, we defined SAP folds as AP folds with a Foldseek probability (FP) ≥ 0.9 and DAP folds as those with an FP < 0.1 to the best structural alignment hit, which was the hit with the highest FP and TM score, in the human proteome unless otherwise specified.

**Fig. 1:**
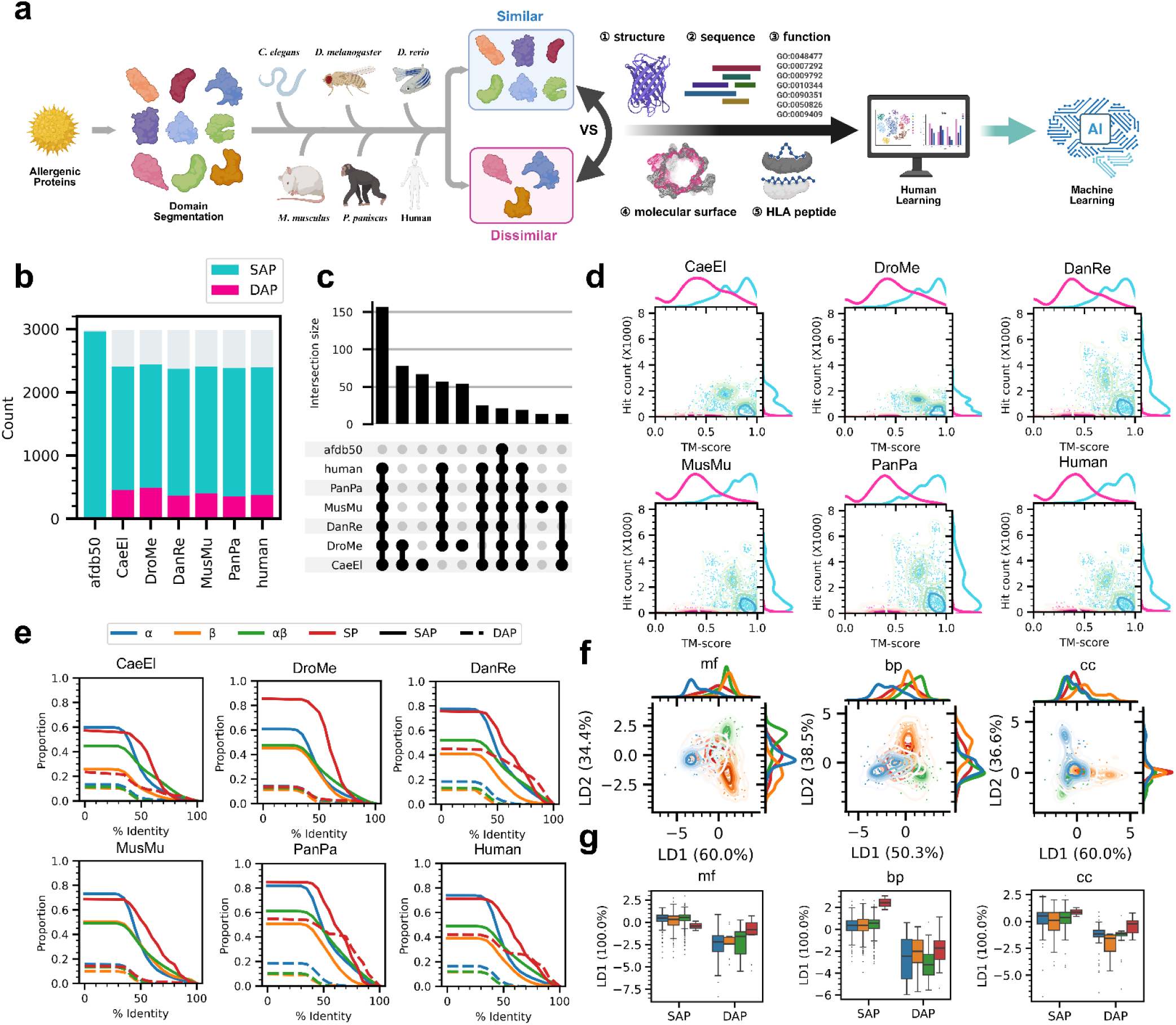
DAP folds map to a distinctive sequence-structure-function space. **a**, Conceptual workflow of this study. **b**, Counts of SAP and DAP folds in afdb50, animal models, and human proteomes. AP folds coloured light grey have medium structural similarity with 0.1 ≤ FP < 0.9. **c**, The most populated intersecting sets of DAP folds with respect to afdb50 and the proteomes. DAP folds with respect to humans were always shared among the mammalian species among these sets. **d**, Distributions of the TM score of the best structural alignment hits (x-axis) and total hit counts (y-axis) in the animal models and human proteomes using SAP and DAP folds with respect to the human proteome as the queries. The 1-D distribution curves shown on both axes were scaled to match the maximum height between SAP and DAP for visibility; thus, the area under the plots should not be interpreted. SAP and DAP colours follow the legend in panel b. **e**, Proportion of fully aligned proteome 20-mers surpassing a threshold of percent identity of alignment to the polypeptide segments of SAP and DAP folds. The proportion at 1.0 represents the total number of alignments, which included 20-mers with no hits and assigned 0% identity, observed in a unique combination of SAP/DAP and CATH fold class. **f**, Functional space comparison supervised for separating the CATH fold classes of the AP folds. CATH fold class colours follow the legend in panel e. **g**, Functional space comparison supervised for separating SAP and DAP folds and shown with CATH fold class division. This projection for the optimal functional segregation between SAP/DAP and among CATH fold classes were apparently incompatible. CATH fold class colours follow the legend in panel e.

On the basis of this definition, we identified unique sequential, structural, and functional features of SAP versus DAP folds (Fig. 1), whose structures are shown in Fig. 2. Most DAP folds were dissimilar to the animal proteomes, prompting us to explore the taxonomic distribution of SAP and DAP folds in afdb50, which revealed strong taxonomical restriction in DAP folds (Fig. 3). We incorporated SNAP folds from food and the human proteome in our comparison, revealing distinct surface features, including B-cell epitope physicochemical profiles, in four ECOD topologies of SAP folds (Fig. 4). These segregating patterns motivated us to develop a machine learning model to classify APs, yielding promising results in predicting unseen APs (Table 1). T-cell epitopes, as important features, led us to elucidate the distinct but shared evolutionary patterns in HLA class II epitopes between SAP and DAP folds (Fig. 5).

**Fig. 2:**
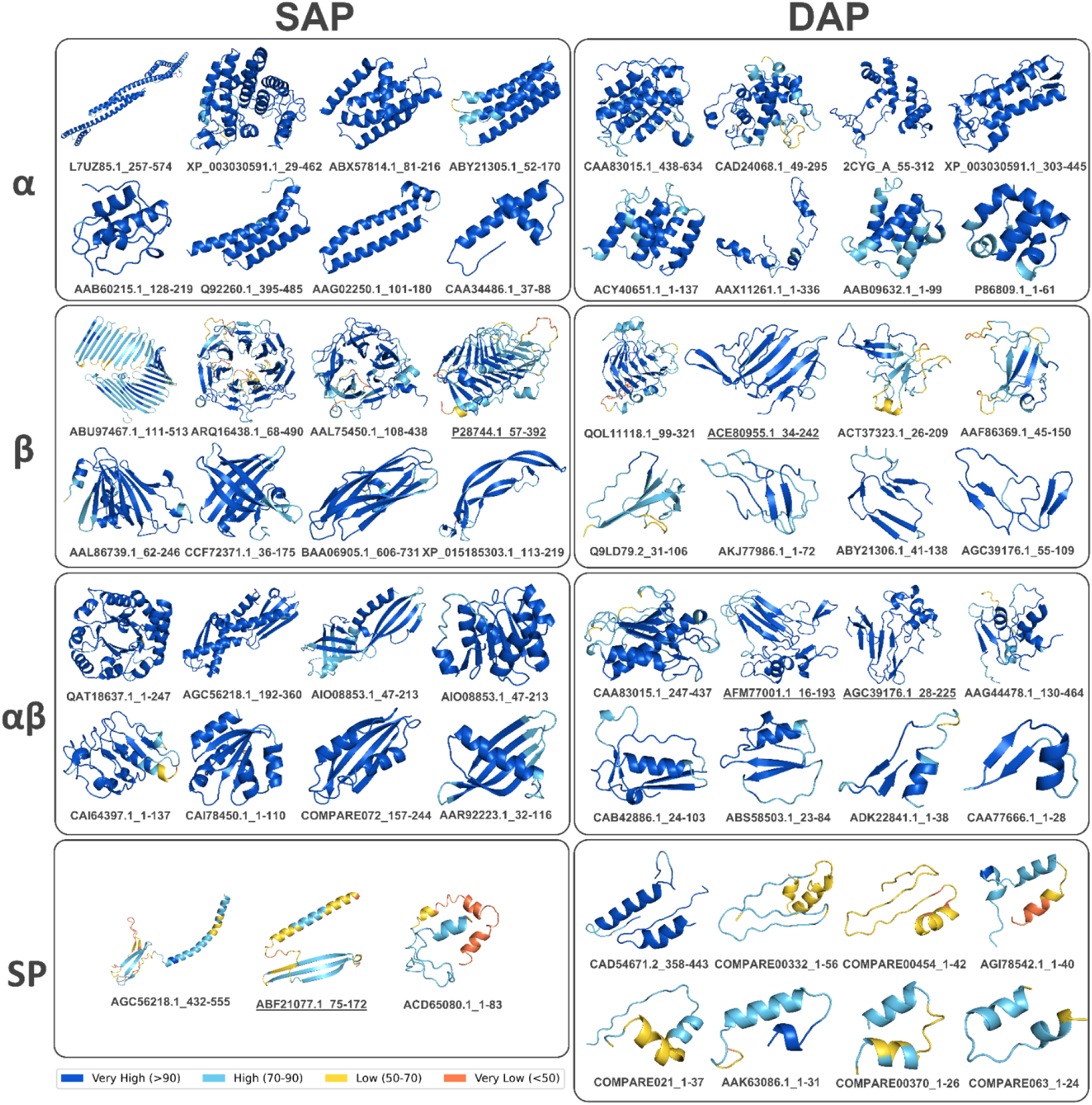
Structures of representative examples of SAP and DAP folds. Structures are coloured according to the pLDDT scores assigned by ESMFold. Structures were sorted in descending order of the number of residues from the top left to the bottom right corner in each specific SAP/DAP-CATH fold class subpanel. Only three representative SAP folds were annotated as the CATH fold class SP. Coexisting SAP-like or DAP-like folds were found in the TED novel fold dataset for the SAP and DAP folds, whose names are underscored. Prefixes of the fold names were the COMPARE database IDs for the original APs. Suffixes of the fold names indicate the start and the end residue position of the AP folds in the original AP sequences.

**Fig. 3:**
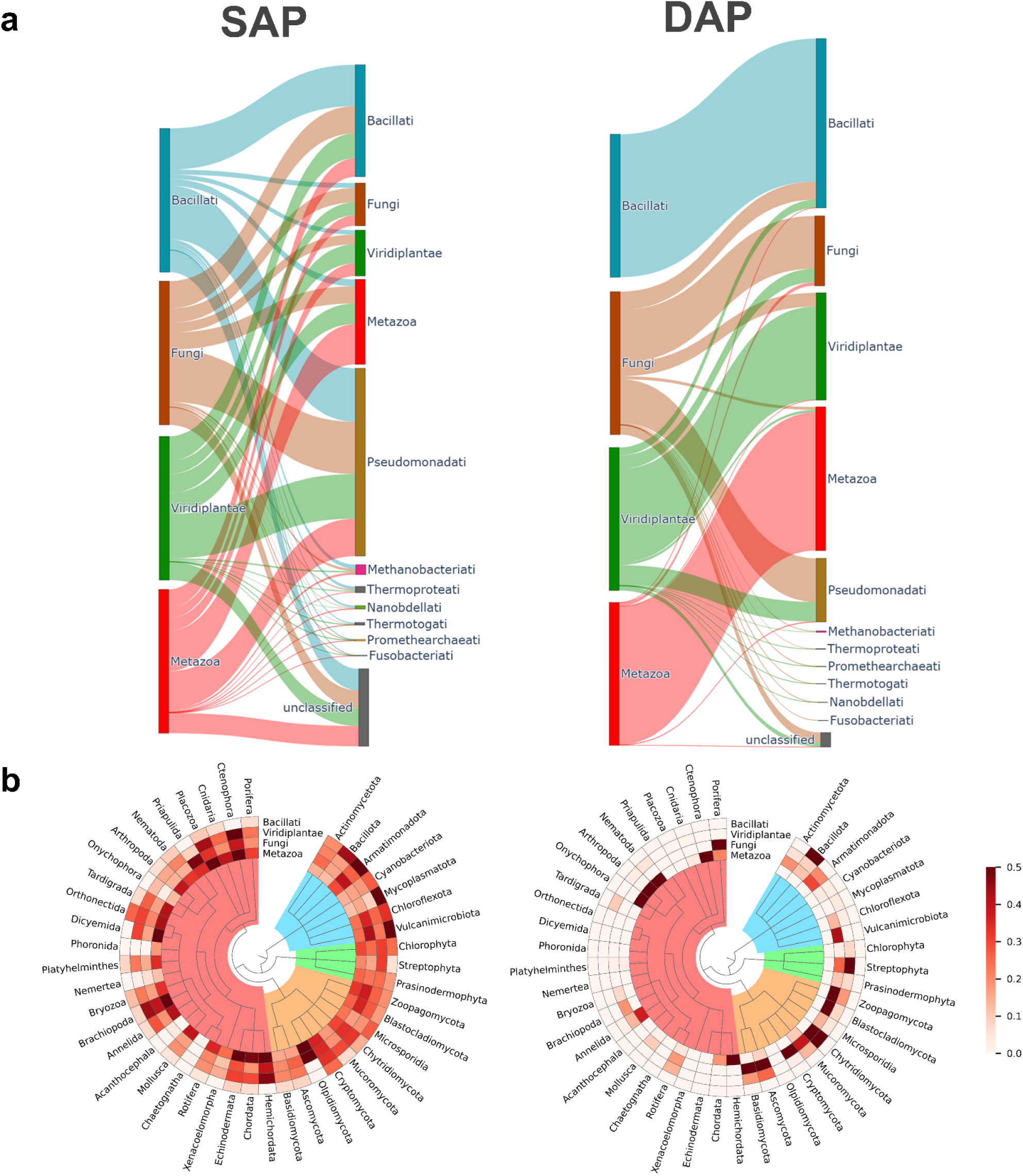
DAP folds are taxonomically restricted. **a**, Kingdom-level Sankey plots visualizing the count of linkages between origin kingdoms of SAP and DAP folds (bars on left sides) and hit kingdoms of SAP-like and DAP-like folds in afdb50 (bars on right sides). The bar sizes of the original kingdoms were standardized. Accordingly, the number of hit kingdoms and the number of unique origin-hit linkages were scaled proportionally. **b**, Phylum-level taxonomic trees overlaid with heatmaps showing the taxonomic organization and abundance of the hit phyla of SAP-like and DAP-like folds in afdb50. Subtrees under metazoa (pink), fungi (orange), viridiplantae (green), and bacillati (blue) are colour shaded. Each circulating channel of the heatmap corresponds to a SAP and DAP fold origin kingdom. The colour of each cell represents the count of origin-hit linkages normalized for over- or underrepresentation of origin kingdoms and the number of existing phyla in afdb50.

**Fig. 4:**
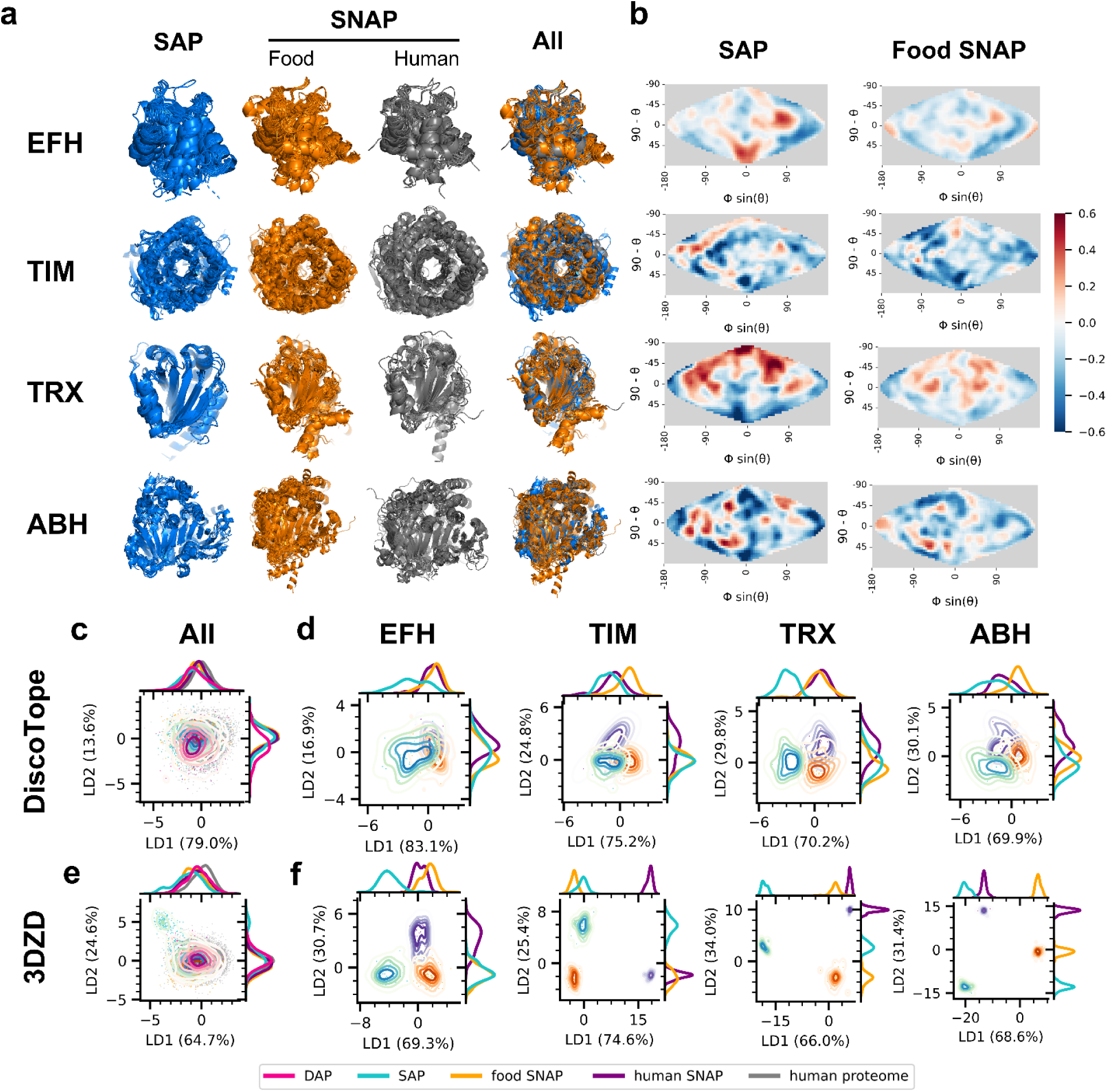
Distinctive surface features of SAP folds compared to food and human SNAP folds. **a**, Structural alignment of the SAP and SNAP folds of the four ECOD topologies. Controlling for their good alignability was a prerequisite for the orientation-specific comparison of surface features. **b**, Projected 2-D maps of the orientation-specific differences in DiscoTope3.0 B-cell epitope scores of SAP and food SNAP from human SNAP folds of the four ECOD topologies. ECOD topology names are shown in rows in Fig. 4a. Blue and red colours indicate lower and higher B-cell epitope scores compared to the human SNAP folds, respectively. Sequence redundancy was reduced to 90%, and resampling by bootstrapping (1,000 iterations) was applied to account for the effect of surface features redundancy and the population size differences among fold classes. **c, d**, Physicochemical space comparison of sized B-cell epitopes supervised for class separation among (c) SAP, DAP, food SNAP, human SNAP, and human proteome folds and (d) SAP, food SNAP, and human SNAP within the same ECOD topology. Changes in the DAP folds were relatively observable in the projected dimension LD2. For each ECOD topology, the presented projection for optimal class separation by LDA generally showed that SAP clusters were more distant than food SNAP clusters from the human SNAP clusters. **e, f**, 3DZD space comparison of the entire surface among (e) SAP, DAP, food SNAP, human SNAP, and human proteome folds and (f) SAP, food SNAP, and human SNAP within the same ECOD topology.

**Fig. 5:**
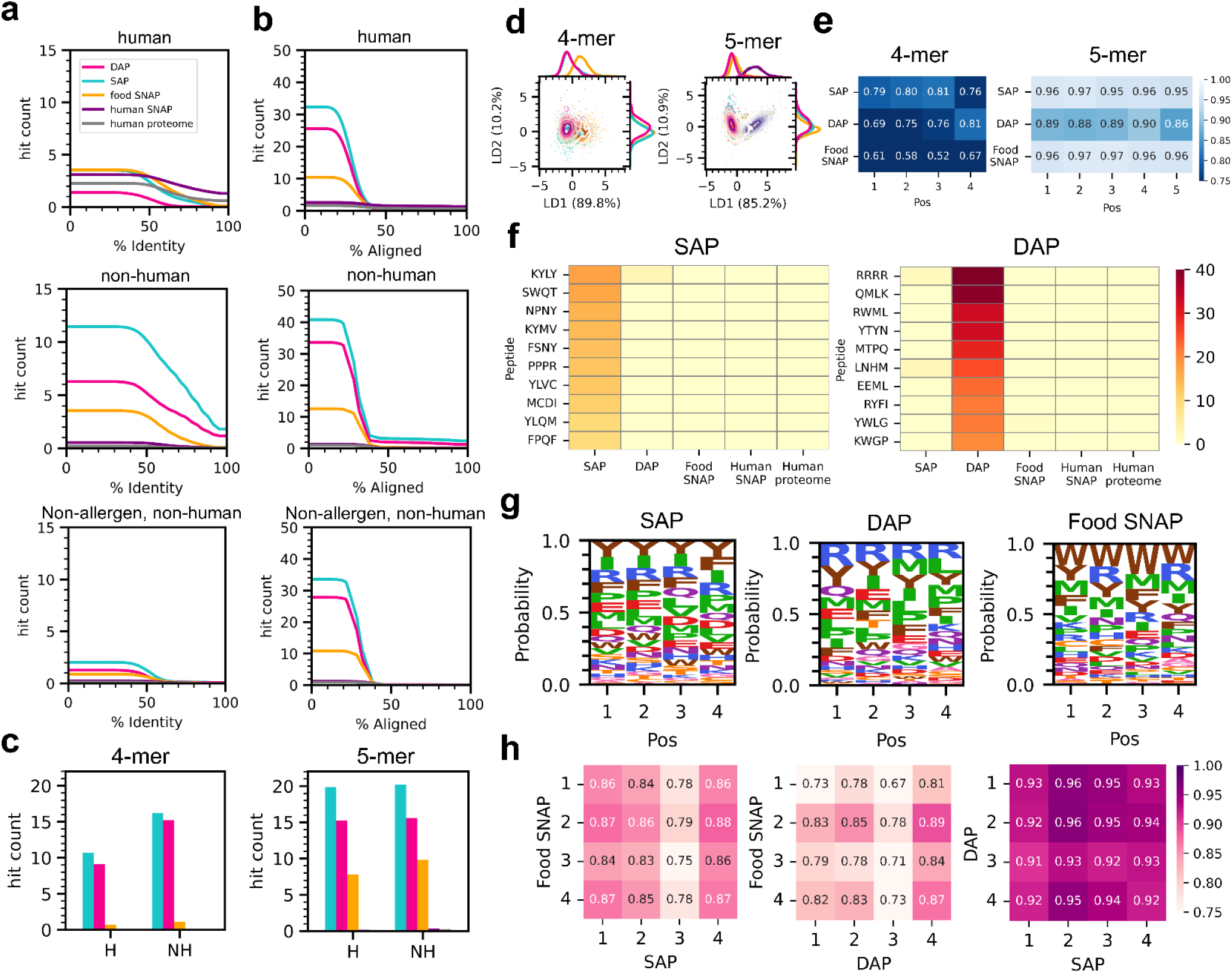
Shared T-cell epitope patterns between SAP and DAP folds. **a, b**, Cumulative hit counts of HLA class II peptides at (a) >90% alignment length and (b) >90% identity of alignment against the polypeptide segments of different fold classes while varying the alternative alignment metric on the x-axis. Hit counts were normalized to the total length of HLA peptides from each source and the total length of the sequence of each fold class. A total of 1,000 bins in the x-axis were used for counting, and rolling averages calculated upon ±5% bins centred at the x-values were plotted for smoothening irregularities due to the cut-off artefacts. **c**, Number of exact matching 4- and 5-mers derived from HLA class II peptides from human (H) and nonhuman (NH) sources. The y-axis was normalized by the same method as in panels a and b. **d**, Sequence space comparison of the HLA class II 4- and 5-mers supervised for class separation among SAP, DAP, and food SNAP by LDA on esm2 embeddings. **e**, Positionwise cos similarity of amino acid frequency from the UniProtKB background among the top 10% most abundant 4- and 5-mers. A lower cos similarity value indicates a more dissimilar amino acid frequency from the background. **f**, Counts of the most abundant, exact matching 4-mers derived from the hit HLA class II peptides in SAP and DAP folds. The counts were normalized by the same method as in panels a and b, and scaled by a factor of 10^11^ for readability. **g**, Sequence logos of the top 10% most abundant 4-mers in terms of SAP, DAP, and food SNAP. **h**, Cross-position cos similarity comparison of the amino acid frequency of the 10% most abundant 4-mers among SAP, DAP, and food SNAP folds.

**Table 1.** The performance of AllerX in the independent test sets, which consisted of the newly deposited APs from COMPARE 2024 and 2025 and a group of APs with structurally similar NAPs in the Maurer Stroh 2019 dataset.

| Dataset | N | Metric | AllerX | AllerCatPro2 | AllerTOPv2.1 | DeepAlgPro |
| --- | --- | --- | --- | --- | --- | --- |
| COMPARE 2024 |  |  |  |  |  |  |
| C24H | 95 | Recall | 0.989 | 0.681 | 0.617 | 0.989 |
| C24N | 46 | Recall | 0.978 | 0.065 | 0.478 | 0.978 |
| All | 141 | Recall | 0.986 | 0.479 | 0.571 | 0.986 |
| COMPARE 2025 |  |  |  |  |  |  |
| C25H | 50 | Recall | 1.000 | 0.600 | 0.580 | 0.940 |
| C25N | 23 | Recall | 1.000 | 0.000 | 0.261 | 0.870 |
| All | 73 | Recall | 1.000 | 0.411 | 0.479 | 0.918 |
| Maurer Stroh 2019 | 386 | Accuracy | 0.484 | 0.857 | 0.750 | 0.607 |
|  |  | Precision | 0.484 | 0.772 | 0.706 | 0.557 |
|  |  | Recall | 1.000 | 1.000 | 0.828 | 0.925 |
|  |  | Specificity | 0.000 | 0.722 | 0.677 | 0.308 |
|  |  | F1 | 0.653 | 0.871 | 0.762 | 0.695 |
|  |  | Roc_auc | 0.500 | 0.861 | 0.752 | 0.616 |
|  |  | Prauc | 0.484 | 0.772 | 0.668 | 0.551 |

Notes:
C24H/C25H and C24N/C25N are the subsets of APs with detectable sequence alignment hits (H) and unseen APs with no detectable sequence alignment hits (N) in COMPARE 2023, respectively.

## Results and Discussion

### DAP folds are rare in evolution

To validate our definition of SAP and DAP folds, we performed the same structural alignment of the AP folds against afdb50, the clustered AlphaFold Protein Structure Database(*24*), as a control. We reasoned that a majority of the DAP folds should have a structural analogue in afdb50 because it should have covered the original proteomes of APs or their representative lineages. Obtaining a clean background in afdb50 was initially challenging because of significant contamination from nonglobular regions, which are common among predicted protein structures(*29*) even after domain segmentation(*30*). We discarded the nonglobular allergenic folds because their “structures” carry little physical meanings; thus, we retained 88.5% of the total number of residues of the segmented AP folds as the globular regions being studied throughout this work, which set a clean background in afdb50 (Fig. 1b) for further analysis.

Our next goal was to trace the gains and losses of SAP and DAP folds in the animal proteomes, inspired by the observed over- and underrepresentation of specific domains in the human proteome compared with those in animal models(*31*). Our structural alignment was sensitive enough to detect these events. For instance, a defensin-like fold (AAO24900.1, 28-82 aa) of the Mugwort pollen allergen Art v 1 was a DAP fold in humans but not in *C. elegans* or *D. melanogaster* (Figs. S1b-e), and the albumin fold was generally a SAP fold in vertebrates but not invertebrates(*32*) (Fig. S1f). However, with respect to the long evolutionary context, we revealed that the number and identity of DAP folds among animal and human proteomes are stable and consistent overall (Figs. 1b,c and S2a-c). We estimated that 67.5% and 12.6% of the AP folds in the COMPARE 2023 database were SAP and DAP folds, respectively (Fig. 1b, Suppl. info, link to “SAP_DAP_count_perc_df.csv”). After structural clustering, the DAP folds were slightly enriched, constituting 22.6% of all the representative AP folds (Fig. S2a). This representative DAP fold percentage was within a ±3% bound, which indicated the relatively stable number of DAP folds among all the animal proteomes. DAP folds were generally underrepresented and structurally different from their best structural alignment hits in the animal proteomes (Figs. 1d and S2d). Similar underrepresentation was also observed in afdb50, although a majority of the DAP folds had at least one structurally similar hit (TM score > 0.5) in the database (Fig. S2d). Collectively, we revealed that DAP folds are underrepresented in evolution and that the gain or complete loss of unique DAP-like folds in animal proteomes was likely an evolutionarily rare event.

Identifying sequence features is crucial to understanding how AP folds might contribute to sensitizing allergic reactions, as molecular recognition relies on specific physicochemical contexts. Contrary to our expectation that SAP folds might have more distinctive sequential features than DAP folds, we found that DAP folds had more dissimilar sequences than SAP folds when compared to the animal and human proteomes, regardless of their CATH fold classes (Fig. 1e). Following the classical sequence–structure relationship, DAP sequences and folds are both more distinctive than SAP folds are, highlighting their strong uniqueness compared with human proteins. With respect to function, we compared the functional space between the SAP and DAP folds. As a control, we identified the distinctive functional space occupied by α, β, and αβ AP folds for molecular function (mf) and biological process (bp), which was reported in general proteins(*33*) but is less obvious in terms of their localization as a cellular component (cc) (Fig. 1f). Using the same approach, we revealed that compared with SAP folds, DAP folds occupied a distinct functional space (Fig. 1g), which was consistent with their localization to a distinct sequence–structure space. For instance, the DAP fold (AAU21501.1, 43–149 aa) of peanut oleosin Ara h 15 was assigned unique functions: “seed oilbody biogenesis” (GO:0010344)(*34*) and “response to freezing” (GO:0050826)(*35*). Another DAP fold (AAB42069.1, 26–114 aa) of the tomato lipid transfer protein Sola l 3 was assigned unique functions, such as calmodulin binding (GO:0005516)(*36*). Similar functions of their homologues were referenced after the GO terms, which were all absent among the SAP folds (Suppl. info, link to “NetGO_GO-Score-min-0.7_GO-centric-summary_LDA-struc-sim.csv”). Our k-nearest neighbour (kNN) distance analysis further elucidated the spatial organization of the SAP and DAP folds within the functional space. We first identified the higher mf and bp functional diversities of αβ AP folds (Fig. S3a), which were consistent with previous observations on the high functional diversity of general αβ proteins(*37*). We demonstrated that compared with the SAP folds, the DAP folds exhibited higher mf functional diversity and potentially slightly restricted cc diversity (Fig. S3b).

Visual comparison of the SAP and DAP folds revealed a striking difference: the DAP folds generally lacked visually appealing fold symmetry, which was more frequent among the SAP folds (Fig. 2). For instance, the SAP folds of the house dust mite allergen α-actinin (L7UZ85.1, 257–537/550–574 aa) and the plane tree pollen allergen Pla or 1 (ABY21305.1, 52–170 aa) were 3- and 4-helix bundles, respectively. In the β and αβ CATH fold classes, multiple SAP folds were symmetrical repeat proteins, such as the 7-bladed β-propeller and the TIM-barrel. In contrast, symmetrical proteins were largely absent among the DAP folds. Notably, some novel SAP-like and DAP-like folds (FP > 0.9 to SAP or DAP folds) coexisted in the TED novel fold dataset(*38*) (Figs. 2 and S4a), highlighting that some APs harbour novel folds that have yet to be experimentally determined.

Additionally, we visually observed the overrepresentation of loop structures in the DAP folds compared with the SAP folds. To this end, we revealed that the random coil content of the DAP folds was slightly higher than that of the SAP folds (Fig. S4b), which could partially contribute to the disrupted fold symmetry and the expected fold instability in the DAP folds. Indeed, we showed that the conformation of the DAP folds was generally less stable than that of the SAP folds (Fig. S4b). A stable fold increases allergenicity through the maintenance of conformational IgE epitopes and resistance to degradation(*39*). Future studies should explore whether DAP folds, in contrast to SAP folds, underlie the differences in protein allergenic potential from a fold stability perspective.

### DAP folds are taxonomically restricted

The absence of most DAP folds in animal proteomes and their underrepresentation in afdb50 imply that animal proteomes are representative subsamples of the known protein universe for the presence or absence of specific allergen folds. This surprising consistency may be attributable to our rigorous definition of DAP folds. Considering the long evolutionary timeframe spanned by the animal models and the stable number of DAP folds among them, we decided to investigate the taxonomic distribution of SAP- and DAP-like folds throughout evolution.

By analysing the taxonomy of the structural alignment hits of SAP and DAP folds in afdb50, we revealed that while SAP-like folds were widespread, DAP-like folds were taxonomically restricted at both the kingdom (Figs. 3a and S5) and phylum levels (Fig. 3b). Notably, bacterial DAP-like folds were found strictly within their kingdom. Similarly, most animal DAP-like folds were found within the kingdom. In contrast, fungal DAP-like folds, followed by plant DAP-like folds, were the most widespread among the four AP origin kingdoms, indicating that although the majority of structural analogues were found within their kingdoms (Fig. S5b), some coexisted in a third kingdom, such as in bacteria and animals (Figs. 3a and S5b). Furthermore, following within-kingdom matches, fungi and plants shared the second-highest number of common DAP-like folds with each other (Fig. S5b). Potential hypotheses for cross-kingdom DAP fold sharing despite their distinctive sequence and folding include horizontal gene transfer (HGT)(*40*), attributed to shared ecological pressures between plants and fungi, or the ancient origin of a minority of these fungal and plant DAP folds, which is supported by their coexistence in archaeal lineages (Fig. 3a). Here, combining the unique sequence features of DAP folds, we propose that cross-reactivity from bacterial and animal DAP folds most likely emerges from AP sources within the same kingdom, followed by fungal and plant DAP folds, whereas the inference for cross-reactive SAP folds is less definitive.

Additionally, we identified Fungi-Porifera and Plant-Chytridiomycota as two notable cross-kingdom, phylum-level linkages for the coexistence of similar DAP-like folds (Fig. 3b). For instance, the DAP fold of the fungus *Aspergillus niger* allergen Asp n 14 (CAB06417.1, 57-112 aa) coexists in the sea sponge species *Amphimedon queenslandica*, which is an early-branching animal, where HGT from alternative kingdoms potentially played a role in its evolution(*41*). Interestingly, multiple DAP folds from fruit allergens such as cherry Pru av 2 (AAB38064.1, 139–227 aa), apple Mal d 2 (AAC36740.1, 158–218 aa), and avocado Pers a 1 (CAB01591.1, 162–219 aa) were found in the fungal phylum Chytridiomycota, an early-branching fungal lineage with significant HGT events(*42*). Chytridiomycota was previously detected in dust samples that were potentially linked to respiratory allergies in children(*43*).

### Unique surface features of SAP folds

Thus far, we have identified distinctive molecular and evolutionary features of DAP folds. An equally important question arises for SAP folds: How do they differ from their nonallergenic structural analogues? To address this question, we carefully investigated the molecular surface of the SAP folds, given its direct relevance to B-cell recognition and the opportunity for a universal analysis of known SAP folds—in contrast to prior studies centred on specific allergen classes(*13*, *15*, *16*, *19*). Accordingly, we compared the surface features of SAP folds against those of SNAP folds across four prevalent ECOD topologies in our AP fold dataset: (1) EF-hand (EFH, 108.1.1), (2) TIM barrels (TIM, 2002.1.1), (3) thioredoxin-like (TRX, 2485.1.1), and (4) α/β-hydrolases (ABH, 7579.1.1). Previous studies often compared allergens to a single group of nonallergens. We reasoned that this approach could yield incidental findings. Therefore, we incorporated both nonself (food) and self (human) SNAP folds in our comparison to reveal surface features that are truly unique to SAP folds.

We mapped B-cell epitope scores onto the surfaces of prealigned SAP and SNAP folds (Fig. 4a and Suppl. info, link to “SAP_SNAP_superpose_count.csv”) and visualized orientation-specific differences in these scores between fold class pairs, namely, (1) SAP versus human SNAP and (2) food SNAP versus human SNAP. We showed that B-cell epitope hotspots were consistently located in similar regions of these SAP and food SNAP folds (Fig. 4b). However, these B-cell epitope hotspots were more pronounced in SAP than food SNAP folds, relative to human SNAP, for EFH, TRX, and ABH topologies, but not for TIM. These findings demonstrate that SAP folds may trigger broader IgE-mediated hypersensitivity compared to food SNAP folds. They also highlight the need for precise SAP-SNAP epitope sorting to prevent hypersensitivity to food SNAP folds.

Extending this analysis, we investigated the physicochemical profiles of sized B-cell epitopes (>600 Å^2^)(*44*), which are more relevant to real-world IgE binding. DAP folds displayed an upshifted global B-cell epitope propensity in these sized B-cell epitopes (Fig. S6a) and a shifted epitope physicochemical profile, whereas SAP, food SNAP and human SNAP folds remained relatively similar (Fig. 4c). Focusing on each ECOD topology, we identified distinct physicochemical profiles for the epitopes across all four ECOD topologies (Fig. 4d). Measurements of intercluster distances in the projected physicochemical spaces further revealed that, relative to human SNAP folds, B-cell epitopes on SAP folds were more distinct than those on food SNAP for EFH, TRX and ABH but not for TIM (Fig. S6b). Among the features driving SAP fold segregation, we identified several Kidera factors(*45*)—KF6 (partial specific volume), KF9 (pK-C), and KF10 (hydrophobicity)—as key alterations in epitope physicochemical properties relative to SNAP folds (Fig. S6c). Consistently, our visualization revealed more pronounced hydrophobic and sticky hotspots on SAP folds than on food SNAP folds (Fig. S7).

We cross-validated SAP fold segregation using rotationally invariant 3D Zernike descriptors (3DZD). The shift in the surface features of DAP folds was apparently not as obvious with 3DZD (Fig. 4e) as with B-cell epitope physicochemical properties at this global scale (Fig. 4c). However, locally within the same ECOD topology, 3DZD allowed clearer separation of SAP than food SNAP from human SNAP folds (Figs. 4f and S6b). This finding suggested that B-cell epitopes alone did not fully account for the molecular surface differences between SAP and SNAP folds, which was consistent with the greater segregation of multiple physicochemical features across the entire surface of SAP folds than food SNAP folds relative to human SNAP folds (Fig. S7).

### Generalizable prediction for unseen APs

Inspired by the distinct molecular signatures of SAP, DAP and SNAP folds, we developed AllerX, a machine learning classifier, to distinguish APs from nonallergenic proteins (NAPs). We integrated sequence, structure, function, molecular surface, and T-cell epitope features, the last due to their critical role in allergy sensitization via HLA class II peptide presentation, which activates Th2 responses. AllerX achieved exceptional performance in the internal test set, correctly classifying 467 of 469 proteins at 98.4% precision for APs from the COMPARE 2023 database (Suppl. info, link to “internal_test.csv”).

To benchmark AllerX, we evaluated its performance against competing classifiers using APs newly deposited in the COMPARE database (versions 2024 and 2025). AllerX excelled in recalling these new APs, including seen or otherwise unseen APs—those without detectable sequence similarity to the training set (COMPARE 2023), matching the strong performance of DeepAlgPro in COMPARE 2024 while surpassing it in COMPARE 2025 (Table 1). In contrast, AllerCatPro2 and AllerTOPv2.1 struggled to identify new APs. In addition, AllerCatPro2 outperformed the others in the Maurer Stroh 2019 dataset, which includes APs and NAPs with similar structures, whereas DeepAlgPro and AllerX showed reduced precision.

To understand AllerX’s ability to detect unseen APs, we compared the importance among features. HLA class II epitope features, expressed as the summed bit score of HLA class II peptide alignments, consistently ranked as the top four important features for the prediction of the internal test and the seen APs from COMPARE 2024 and 2025 (Fig. S8a). These features, which quantify the presence of HLA class II epitopes, effectively distinguished NAPs from the training set APs, as evidenced by their segregated distributions (Fig. S8b). However, for unseen APs, these bit score distributions were even lower than those of NAPs, reflecting the underrepresentation of known HLA class II peptides in their sequences, potentially owing to their novelty, which poses a challenge for accurate classification by AllerX. In response, AllerX reprioritized a sequence (esm15) and a structural (progres_063) feature, both of which were ranked ≤ 4, exclusively for unseen APs (Fig. S8a), which negatively correlated with the allergen class (r^2^=0.44 and 0.52, respectively) but were not intercorrelated (Fig. S8c). Moreover, the ranks of the two nonhuman-origin HLA class II features decreased slightly (Fig. S8a). Combined with the highly ranked (≤ 3), human-origin HLA class II features, the sequence and structural features aided AllerX’s robust prediction of unseen APs.

### Shared patterns in HLA class II 4-mers

Guided by the feature importance analysis in AllerX, we further investigated how the quantity and quality of HLA class II epitopes differ among SAP, DAP, SNAP, and human proteome folds. We compared the number of sequence alignment hits of HLA class II peptides in their sequences by imposing a stringent filter of 90% alignment length of the peptides while varying the percent identity of alignment (Fig. 5a) and flipping the metrics (Fig. 5b). As expected, we showed the overrepresentation of allergen source HLA class II peptides at a high percentage alignment in SAP and DAP, followed by a medium percentage in food SNAP, and the underrepresentation in human proteins (Fig. 5a). We compared these results using T-cell epitope assignment by NetMHCIIpan, which revealed the overrepresentation of HLA class II epitopes for both the DR and DQ allele subtypes in SAP folds but not for DR in DAP folds (Fig. S9a). Unexpectedly, we detected alignments at high percent identity of truncated (30–40% full-length) HLA class II peptides from both human and nonhuman sources, including those from nonallergens (Fig. 5b). We revealed that these nonallergen peptide sources were mostly pathogenic viruses and bacteria (Suppl. info, link to “blast_df_selected2_copy_non-allergen_non-human_II_pident90.csv”). Relatedly, the sequence and structural similarities between allergenic and parasitic proteins have been previously reported(*13*). Here, we demonstrated the linkage between APs and pathogenic viruses and bacterial proteins in terms of truncated segments of their T-cell epitopes.

Next, we demonstrated that a majority of the truncated HLA class II peptides composed a group of 4- and 5-mers with exact matches in SAP and DAP sequences (Fig. 5c) and a minority of 6-mers (Fig. S9b). Despite the exact matches, the very short alignment lengths prompted us to assess whether the HLA class II k-mers encoded any biological information rather than mere random noise. To this end, we first demonstrated that compared with those from food SNAP folds, 4- and 5-mers from SAP folds and DAP folds localized to a more proximal sequence space (Fig. 5d), although the result was less conclusive for 6-mers (Fig. S9c). Second, compared with those of the 5- and 6-mers, the positionwise amino acid frequencies of the 4-mers were more dissimilar from the background amino acid propensity (Figs. 5e and S9d, e), highlighting the potential biological information presented by the 4-mers compared with the 5- and 6-mers.

We then isolated the most abundant 4-mers in the SAP and DAP folds. Intriguingly, these abundant 4-mers were almost exclusively found within their own SAP or DAP fold class (Figs. 5f and S9f), and many of them were confined to a limited number of AP groups; for instance, the SAP fold 4-mer KYLY was shared in the *Penicillium* serine protease Pen ch 18 (AAF71379.1, 33-472 aa) and the boar albumin Sus s 1 (NP_001005208, 28-226 aa), and the DAP fold 4-mer QMLK was shared in plant chitinases such as the banana Mus a 2 (CAC81811.1, 67-155/213-311 aa) and avocado Pers a 1 (CAB01591.1, 75-161/220-326 aa) (Suppl. info, link to “456mer_blast_df_selected5_logo.csv”). We thus further compared the sequence conservation of 4-mers among SAP, DAP, and food SNAP folds (Fig. 5g). Consistent with the sequence space analysis, compared with those in food SNAP folds, the 4-mers in SAP and DAP folds were more similar in terms of amino acid frequency across positions (Fig. 5h). While the sequence and the source of these SAP and DAP 4-mers were distinct, they surrounded a closer sequence space than those of food SNAP folds did, even those that constructed similar folds as SAP folds but not DAP folds. This finding demonstrated a fold-independent, sequence–space linkage between the T-cell epitopes of SAP and DAP folds.

Although HLA class II 3-mers could be present for T-cell activation(*46*), whether these 4-mers alone could form a stable antigen-presenting complex with HLA class II molecules is unknown. Considering the readily comparable amino acid frequencies across positions (Fig. 5g), we estimated the likelihood of these 4-mers binding to HLA-DR and HLA-DQ when primed at the P1-P6 positions in both the N➔C and the alternative C➔N orientations(*47*), which revealed a generally greater likelihood of binding to both allele types with the 4-mers from the SAP and DAP folds than to those of the SNAP and human proteome folds (Fig. S10). Additionally, the functions of some of these SAP-fold 4-mers have been previously reported. For instance, upon tyrosine phosphorylation, the SAP 4-mer NPNY could serve as a docking site for PTB domain-containing proteins such as SHC, thereby activating mast cell degranulation via the PI3K/AKT and MAPK pathways(*48*, *49*). Another SAP 4-mer, KYMV, is a subsequence of the bioactive peptide WKYMVM, which can stimulate PIP2 hydrolysis into DAG, activating PKC to promote B-cell activation(*50*). These findings echoed the alternative functional roles of some of these 4-mers, other than direct T-cell activation through antigen presentation.

## Conclusion

Our comparative analysis of SAP, DAP, and SNAP folds establishes a dual-AP schema, offering key insights into the host–factor evolution of allergenic responses. DAP folds are evolutionarily underrepresented, taxonomically restricted, and highly dissimilar to human proteins in sequence and structure. Consequently, immune encounters with DAP folds are likely sporadic. Their distinctive molecular fingerprints would have been easily perceived as foreign throughout much of the evolution of the human immune system. The observation that fungal and plant DAP-like folds are prevalent across kingdoms, whereas bacterial and animal DAP-like folds are kingdom-specific, enhances source-tracing capabilities for cross-reactive APs in food safety incidents.

Conversely, SAP-like folds, as ubiquitous environmental “noise”, drive continuous SAP–SNAP sorting by the immune system. The distinct molecular features of SAP folds, particularly B-cell epitopes, exhibit specific physicochemical profiles, supporting intrinsic allergenic determinants and challenging arbitrary notions of allergenicity. Host factors, such as cytokine profiles, likely coevolved with the ubiquity of SAP folds, promoting balanced Th1/Th2 immune skewing to prevent autoimmunity while maintaining surveillance against AP-like pathogen proteins. This contrast between the continuous immune engagement of SAP folds and sporadic encounters with DAP folds likely suggests a dual-path host–factor evolution in allergenic responses to SAP and DAP folds. Identifying common B-cell epitopes on SAP folds and the shared T-cell epitope patterns between SAP and DAP folds facilitates the rational design of broad-spectrum allergy vaccines. Our classifier, AllerX, shows promising predictive performance for unseen APs, validating our multifaceted approach to detect APs for improved food safety.

## Methods

### Domain segmentation and alignment

AP sequences were downloaded from the COMPARE database(*51*) on 2023 Nov. AP structures, except those that contained ambiguous amino acid residues, were predicted by ESMFold(*52*), yielding 2,226 AP structures. AP folds were segmented by Merizo(*30*), yielding 3,496 AP folds. The residualwise globularity of each AP fold was annotated by AlphaCutter(*29*). An AP fold was considered nonglobular and discarded if all of its residues were annotated as nonglobular, retaining 2,985 AP folds. Protein structures of the animal proteomes of *C. elegans, D. melanogaster, D. rerio, M. musculus, P. paniscus,* and humans were obtained from the AlphaFold Protein Structure Database(*24*) and segmented into domains according to the methods of Schaeffer et al.(*31*) Each AP fold was structurally aligned against the animal proteome domains and afdb50 using Foldseek(*26*) (version 8-ef4e960, downloaded Dec 1 2023) with the option --max-seqs = 1,000,000 to ensure that all hits were reported. A total of 880 representative AP folds were annotated by structural clustering using Foldseek easy-cluster with an alignment coverage = 0.9, from which example structures are shown in Fig. 2.

### Food and human SNAP fold selection

For food SNAP folds, first, UniProt protein IDs were collected from the AlphaFold Protein Structure Database(*24*), selecting species that are major food allergen sources(*53*) with NCBI taxonomy IDs: 3847 (soybean), 4565 (wheat), 4182 (sesame), 3818 (peanut), 8030 (salmon), 8236 (tuna), 27697 (sardine), 41210 (crab), 6706 (lobster), 6687 (shrimp), 3755 (almond), 51240 (walnut) and 13451 (hazelnut). There were two reasons to select SNAP folds from these food allergen sources. First, it established a comparable evolutionary context to that of APs to minimize the effects of this context on the molecular characteristics of SNAP folds. Second, real-world immune encounters with standalone SNAP folds were justified by food protein digestion and absorption. These food protein UniProt IDs were further filtered by (1) a list of the same Foldseek cluster representative IDs(*54*) as the best structural alignment hits of APs in the human proteome and (2) ensuring that they were not flagged with “Allergenic Properties” in UniProt. In parallel to the generation of the AP fold dataset, these food proteins underwent the same structure prediction, domain segmentation, and structural alignment against human proteome folds, using the same FP cutoff > 0.9 for the best structural alignment hits. These food SNAP folds, were used in the B-cell epitope physicochemical space analyses in Figs, 4c and e, and the T-cell epitope analyses in Fig. 5. For the surface analyses for specific ECOD topologies in Figs. 4a, b, d, and f, the food SNAP folds were further selected according to the following sequence length criteria: (1) alignment length to the best structural alignment hit in the human proteome fold < 10% difference for both the length of the food SNAP fold and the best hit, and (2) a food SNAP fold sequence length < 2 SD from the mean length of the human proteome folds with the same ECOD topology ID. These length criteria were applied to ensure the absence of major insertions or deletions in the food SNAP folds for good structural alignability; the inclusion of sequences falling outside these length criteria would hinder our surface analyses, which were focused on physicochemical properties instead of fold shape differences. For these length-selected food SNAP folds, we also ensured that at least five of them had a B-cell epitope >600 Å^2^ within their ECOD topology group to ensure a minimal population size for comparison—finally leaving us only the four ECOD topologies EFH, TIM, TRX, and ABH of the SAP folds being studied.

The human SNAP folds used in the B-cell epitope physicochemical space analyses in Figs, 4c and e, and the T-cell epitope analyses in Fig. 5 were the best structural alignment hits in the human proteome folds pooled from the structural alignments from (1) SAP and (2) food SNAP folds before sequence length or B-cell epitope number selection. The human SNAP folds used in the surface analyses in Figs. 4a, b, d, and f were the best structural alignment hits corresponding to the SAP and food SNAP folds after sequence length and B-cell epitope number selection.

### Proteome k-mer alignment

The model animal proteome sequences were downloaded from UniProt using the proteome IDs UP000001940 (*C. elegans*), UP000000803 (*D. melanogaster*), UP000000437 (*D. rerio*), UP000000589 (*M. musculus*), UP000240080 (*P. paniscus*), and UP000005640 (human). Each proteome sequence was divided into all possible 20-mers. BLASTp of BLAST+ 2.15.0(*55*) was used for sequence alignment with the 20-mers as queries to the AP-fold polypeptide segments as alignment subjects with the option -max_target_seqs = 10^9^ to ensure that all hits were reported. Alignments were retained only if the whole 20-mer was aligned. The cumulative numbers of aligned k-mers surpassing an increasing percent identity of alignment were counted, where k-mers without any hits were counted as a virtual alignment with 0% identity.

### Functional latent representation

Polypeptide segment sequences were concatenated for each AP fold, where the sequence redundancies of the concatenated sequences were reduced to 90% by CD-HIT(*56*). NetGO 3.0(*57*) was used to assign GO terms to each AP fold using the representative sequences as input. To minimize assignment artefacts due to sequence concatenation, the GO terms of the full-length APs were also assigned. We retained an assigned GO term in an AP fold only if it also appeared in the assignment to the respective full-length AP. Assigned GO terms were further selected on the basis of a NetGO score > 0.7. AP folds with < 3 assigned GO terms for any GO term type (mf, cc, and bp) were not considered because they introduced a strong bias to the spatial distribution analysis in functional space. Each GO term was embedded by anc2vec(*58*). The means of the anc2vec vectors for each GO term type were used as the latent functional representations of each AP fold. Linear discriminant analysis (LDA) was used for the dimension reduction of the functional representations down to two- and one-component supervised on CATH fold classes and SAP/DAP fold classes, respectively. For the spatial distribution analysis, kNN distances by cosine similarity between the means of anc2vec vectors for k=1–100 were computed for each GO term type. To account for population size differences among CATH fold classes and between SAP/DAP folds, bootstrapping (1,000 iterations) was applied, and proteins were sampled to match the smallest fold class within each GO term type to obtain mean kNN distances. These mean kNN distances were normalized to those of the SAP folds or αβ class at k=100.

### Taxonomic analyses

All structural alignment hits in afdb50 of the representative AP folds were retrieved for their kingdom- and phylum-level NCBI taxonomy. The numbers of unique linkages between the AP-fold origin kingdom (i.e., Bacillati, Fungi, Viridiplantae, and Metazoa) and afdb50-hit kingdom or phylum, denoted as “origin-hit linkage” here, were counted. For the Sankey plot (Fig. 3a), to enable a standardized comparison among AP-fold origin kingdoms, the number of each origin kingdom was resampled to 10,000, while the numbers of origin-hit linkages were scaled accordingly. For the kingdom-level heatmap (Fig. S5), each count of origin-hit linkages was normalized by (1) the total counts of origin-hit linkages of the corresponding AP-fold origin kingdom and (2) the total number of unique kingdoms that existed in afdb50 and then scaled by a factor of 10^7^ for readability. This enabled a standardized comparison among all origin-hit linkages by accounting for over- or underrepresentation of certain AP fold origin kingdoms and the number of existing kingdoms in afdb50. The same normalization and scaling were applied to the phylum-level heatmap (Fig. 3b) but with phylum counts instead of kingdom counts. Taxonomic trees were constructed by phyloT(*59*) with the hit phyla NCBI taxonomy IDs and visualized by pycirclize(*60*).

### Molecular surface analyses

The SNAP fold selection procedure for the molecular surface analyses in Fig. 4 was detailed in the Methods subsection “Food and human SNAP fold selection”. The Relax function of Rosetta (2021.16)(*61*) was used to resample side-chain rotamers of SAP, DAP, SNAP, and human proteome folds to eliminate the differences in side-chain optimization bias between ESMFold and AlphaFold. Folds across all fold classes with the same ECOD topology were structurally aligned to a single consensus alignment using US-align(*62*) with the option -mm = 4. B-cell epitopes were assigned on the folds by DiscoTope3.0(*63*) using all default settings. DsicoTope3.0-calibrated B-cell epitope scores (Fig. 4b) and physicochemical properties on the fold surface (Fig. S7) were visualized by SURFMAP(*64*) using representative folds, which were further reduced to 90% sequence redundancy of the concatenated polypeptide segments by CD-HIT(*56*) of each fold class to control surface features redundancy, as inputs. To calculate the difference in B-cell epitope scores and physicochemical properties between SAP and food SNAP folds to human SNAP folds while accounting for the population size differences among fold classes, bootstrapping (1,000 iterations) was applied, sampling folds to match the smallest fold class within each comparison to obtain mean values at each SURFMAP cell, which corresponded to a specific orientation of the consensus alignment. Differences at each SURFMAP cell were obtained by subtracting the mean values of SAP or food SNAP from those of human SNAP folds.

For the comparison of B-cell epitope physicochemical space (Figs 4c and d), first, B-cell epitopes were isolated by aggregating surface residues that were (1) assigned as a B-cell epitope by DiscoTope3.0(*63*), (2) if they were in contact (any heavy atoms of a pair of residues ≤ 4.5 Å)(*65*), and (3) if the total SASA of the aggregated epitope was >600 Å^2^. The physicochemical properties of each B-cell epitope were assigned as the sum of each 66 aa descriptor(*66*) of the epitope residues. The fold-level physicochemical properties were further calculated by summing the sums of the epitope-level aaDescriptors and then normalizing them to the total SASA of all the B-cell epitopes. LDA, supervised by fold classes based on the normalized sums of the aaDescriptors (Fig. 4c and d) and 3DZD(*67*) (Fig. 4e and f), was used for dimension reduction down to two components. In contrast to the physicochemical space analyses in Figs. 4c and d which involved surface selection for sized B-cell epitopes, 3DZD(*67*) (Fig. 4e and f) embedded the entire fold surfaces. For the B-cell epitope score distribution shown in Fig. S6a, DiscoTope3.0-uncalibrated scores were used because customized normalization was performed by epitope SASA, which accounts for the influence of the epitope size on the score(*63*). Feature importance was analysed by SHAP(*68*) throughout this study.

### AllerX data collection

To collect NAPs for AllerX training, we first identified the best sequence alignment hits in the UniProtKB/Swiss-Prot (downloaded 2024 Oct) of each COMPARE 2023 AP sequence as a query with BLASTP. NAPs for AllerX training were collected from three sources: (1) proximal sequence space: member sequences of the same UniRef50 (N=745) and UniRef90 (N=415) clusters as the best sequence alignment hits; (2) proximal fold space: member sequences of the same Foldseek clusters(*54*) (N=1,869) as the best sequence alignment hits; and (3) random UniProtKB/Swiss-Prot sequences (N=3,804). All the NAP sequences were confirmed as not flagged with “Allergenic Properties” in UniProt. We reasoned that this selection scheme, covering the proximal and distant NAPs in the sequence and fold space, enabled a better representation of NAPs in contrast to similar attempts, which favoured random selection(*69*). These NAPs (N=6,833), as the negative class (0), together with COMPARE 2023 APs (N=2,536) as the positive class (1), formed the AllerX training dataset, which was used for training, k-fold validation, and internal testing. NAP structures from this training set were downloaded from the AlphaFold Protein Structure Database(*24*). NAP and AP structures from the independent test sets, which included COMPARE 2024 (N=141), COMPARE 2025 (N=73), and the Maurer Stroh 2019 dataset(*70*) (N=386), were predicted by Boltz-1(*71*), which was released at the time and was used for accurate protein structure prediction.

### AllerX training and validation

Sequence embedding was extracted from the 6th layer of esm2_t6_8M_UR50D(*52*). Structural embedding was extracted from Progres(*72*). Functional embedding was extracted from anc2vec(*58*) on the basis of GO terms with scores > 0.7 assigned by NetGO 3.0(*57*). Molecular surface was embedded by 3DZD(*67*). For T-cell epitope-related features, first, the IEDB MHC peptide database(*73*) was downloaded. Only records (1) with humans as the host, (2) with the identified epitope source organism, (3) with peptidic epitopes, and (4) without ambiguous amino acids in the sequence were selected. These HLA peptide sequences were further divided into four categories according to the following criteria: (1) human or nonhuman as the epitope source and (2) class I or II as the HLA peptide. Sequence redundancy reduction to 90% by CD-HIT(*56*) was applied to each HLA peptide category, which was then used as a query in BLASTP-short against the training and independent test sequences as the alignment subjects with the option - max_target_seqs = 1,000,000. We filtered the alignments by (1) whether the aligned HLA peptide length was ≥ 90% of its full length or not and (2) whether the percent identity of alignment was ≥ 90% or not and summed their bit score per protein with its normalized length, creating four filtered groups of alignment hits. These four filtered groups of alignment hits, compounded with the four HLA peptide categories, yielded a total of 16 T-cell epitope features that described the alignment quality of HLA peptides from different sources and classes.

Five percent of the records in the AllerX training dataset were split into an internal test set, with the remaining 95% used as the training and validation set. ShapRFECV(*74*), with an XGBoost binary classifier(*75*) trained on the training and validation sets and optimized for the f1 (macro) score, was used for feature selection down to a subset of 83 features (Suppl. info, link to “shap_values_summary.csv”). By 3-fold validation with training:validation split at 8:2 for each iteration, the best model with the highest validation PR-AUC was used for internal testing and independent testing (Table 1).

For the independent test set prediction by the competing AP classifiers, the AllerCatPro2 and AllerTOPv2.1 web servers at all default settings were used for their predictions. The pretrained model weight of DeepAlgPro, deposited on GitHub, was used for prediction using all default settings. The outputs of the AllerTOPv2.1 and DeepAlgPro prediction results were natively binary. For the AllerCatPro2 prediction results, proteins labelled “strong evidence” were annotated as the positive class; otherwise, they were annotated as the negative class, which were assigned the original labels “weak evidence” or “no evidence”.

### T-cell epitope analyses

HLA class II peptides shown in Figs. 5a and b were counted from a separate BLASTP-short alignment with the same setting in the generation of T-cell epitope-related features in AllerX but the alignment subjects were the polypeptide segments of SAP, DAP, SNAP, and human proteome folds. Hit counts were normalized by the total length of HLA peptides from each source and the total length of the sequence of each fold class and then scaled by a factor of 10^8^ for readability. For the T-cell epitope assignment by NetMHCIIpan4.3(*76*), the 90% sequence redundancy-reduced polypeptide segments of the SAP, DAP, SNAP, and human proteome folds were used as the input. Any segments with <10 residues were not considered, which was a constraint of NetMHCIIpan4.3(*76*). A list of HLA class II alleles of the subtypes DR and DQ was curated for their documented association with protein allergen-induced allergy (Suppl. info, link to “HLA_classII_allele_subtype_selection_support_evidence.doc”) and was used for the assignment targets. Alleles from the DP subtype were not included because of a lack of similar documentation. The counts of HLA class II peptides predicted for binding were normalized by the number of curated alleles for each subtype and the total length of polypeptide segments for each fold class and then scaled by a factor of 10^3^ for readability (Fig. S9a). For the sequence space analysis of HLA class II k-mers (Figs. 5d and S9b), all unique and exact matching k-mers in the sequences of SAP, DAP, SNAP, and the human proteome folds were embedded by the 6th layer of esm2_t6_8M_UR50D(*52*), which was then dimensionally reduced to two components supervised on the fold classes using LDA. For the positionwise amino acid frequency similarity matrices (Figs. 5e, h and S9d), cos similarity was used as the distance metric between amino acid frequencies as 20-D vectors. The UniProtKB background amino acid frequency was obtained from the database statistics(*77*).

### HLA class II k-mers binding likelihood

To evaluate the HLA molecule binding potential (Fig. S10), the top 10% most abundant HLA class II k-mers for each fold class were evaluated using PSSM matrices built upon the core HLA peptides (P1-9) from the MHC Motif Atlas(*78*) with the same set of curated DR and DQ alleles, which was used in NetMHCIIpan4.3(*76*) prediction, as targets. A pseudocount of 0.01 was assigned for unobserved amino acids. The PSSM score was defined as the probability of each amino acid in the PSSM at its aligned position (P1-6) and scaled by a factor of 100 for readability. With a given simulated k-mer priming (e.g., a 4-mer primed at P1 in the N➔C direction), the mean PSSM score, which averaged the PSSM scores of the aligned positions (i.e., P1 to P4 as in this example), was used to represent the binding potential of the k-mer.

### CATH fold class classification

All SAP, DAP, SNAP, and human proteome folds were assigned to their CATH fold class with a classifier trained on the CATH40(*79*) data using DSSP(*80*) secondary structure annotations as features and validated with an overall 90.0% classification accuracy across CATH fold classes. The class FS was not considered in this study because no SAP folds were assigned as FS.

### Fold stability analysis

We generated ensembles of AP folds by sampling 1,000 snapshots each using BioEmu(*81*). The C-α RMSD of each snapshot was calculated using the first snapshot as the reference.

### Figure and molecular visualization

Data analysis and visualization were performed using Python v.3.11.7 (https://www.python.org/) with the following libraries: NumPy v.2.0.2 (https://github.com/numpy/numpy), pandas v.2.2.2 (https://github.com/pandas-dev/pandas), Matplotlib v.3.10.0 (https://github.com/matplotlib/matplotlib), seaborn v.0.13.2 (https://github.com/mwaskom/seaborn), Plotly v.5.24.1 (https://github.com/plotly/), scikit-learn v.1.6.1 (https://github.com/scikit-learn/scikit-learn), SHAP v.0.48.0 (https://github.com/slundberg/shap), UpSetPlot v.0.9.0 (https://github.com/jnothman/UpSetPlot), pycirclize v.1.9.1 (https://github.com/moshi4/pyCirclize), Biopython v.1.85 (https://github.com/biopython/biopython), pymsaviz v.0.5.0 (https://github.com/moshi4/pymsaviz), and Google Colaboratory (https://research.google.com/colaboratory). The data in Fig. 1a were created using BioRender (https://www.biorender.com/). Structural visualizations were generated with PyMOL v.2.55.5 (https://github.com/schrodinger/pymol-open-source). Molecular surface visualizations were generated with SURFMAP v.2.2.0 (https://github.com/i2bc/SURFMAP). In Figs. 1d, f, 4c-f, 5d, S2d, and S9c, the distributions expressed as 2-D contours and 1-D curves were plotted by kernel density estimate with seaborn.

## Acknowledgements

The author has no acknowledgements.

## Author contributions

CLT conceived the study, performed the research, analysed the data, and wrote the manuscript.

## Data and materials availability

All AP fold structures and the generated datasets from this study were deposited in https://zenodo.org/records/XXXXXXXXXXXX. Codes used for the data processing and figure generation were deposited in https://github.com/XXXXXXXXXXX. Codes for feature engineering, training, and testing in the AllerX model, and its pretrained model weights were deposited in https://github.com/XXXXX

