## Supplementary material for "Dual schema of allergens reveals molecular and evolutionary signatures": suppl. info

Chunlai Tam[1]

[1]Department of Integrated Biosciences, Graduate School of Frontier Sciences, The University of Tokyo, Tokyo, Chiba, Japan.

**Fig. S1: FP distribution of best structural alignment hits and the loss and gain of AP folds in the human proteome. a,** FP distribution of the best structural alignment hit in afdb50 and animal model proteomes with all AP folds as query. **b-e,** Tertiary, secondary structure, and sequence alignments among the defensin-like fold (AAO24900.1, 28-82 aa) of the Mugwort pollen allergen Art v 1 and its best structural alignment hits in the animal models and human proteomes. **b,** TM score distribution of the best structural alignment hits in the animal models and human proteomes using DAP folds with respect to the human proteome as the query. This analysis of the defensin-like fold was initiated by the observation of a gradually diminishing shoulder region from CaeEl to human at TM score around 0.6 to 0.8, which implied the loss of structural alignability of some DAP folds. **c,** Tertiary structural alignment of the defensin-like fold (white) with the best structural alignment hits in each proteome, which was grouped by protostomes (red), deuterostomes (green), and hominids (cyan). TM score was annotated below the species names. A sudden drop in TM score was observed between deuterostomes and hominids. **d,** Secondary structure alignment based on the tertiary structural alignment. α-helices (purple), β-strands (green), loops (yellow), and the alignment gaps (grey) were annotated. C-terminal insertions were observed in the best hits in deuterostomes and hominids but not in protostomes. **e,** Sequence alignment of the best structural alignment hits based on the tertiary structural alignment. **f,** Albumin fold was general a DAP and a SAP fold in invertebrates (IV) and vertebrates (V), respectively. Albumin folds (white) were aligned with the best structural alignment hits (colored) in the animal models and human proteomes. TM score was annotated below each structural alignment.

**Fig. S2: Abundance of representative and CATH fold-divided SAP and DAP folds. a-c,** Count of representative SAP and DAP folds in afdb50, the animal models, and the human proteomes. **a,** Counts of representative SAP and DAP folds. **b,** CATH fold class-divided counts of representative SAP and DAP folds. **c,** Most populated intersecting sets of representative DAP folds with respect to afdb50 and the proteomes. Compared to the unclustered AP folds (Fig. S1c), there was a stronger contrast between the size of the most populated intersecting DAP set, which shared among all the animal models and the human proteomes, versus unique DAP sets that shared within a limited number of proteomes (e.g. *C. elegans* or *D. melanogaster).* **d,** CATH fold-divided distributions of TM score of the best structural alignment hits (x-axis) and total hit counts (y-axis) in afdb50, the animal models, and the human proteomes using SAP and DAP folds with respect to the human proteome as the queries. The 1-D distribution curves shown on both axes were scaled to match the maximum height between SAP and DAP for visibility; thus, the area under the plots should not be interpreted.

**Fig. S3: Spatial organization in functional space of AP folds divided by (a) CATH fold classes, and (b) SAP and DAP folds**. The spatial organization was expressed by the mean kNN distance (solid line) and the shaded area of ±1 SD bounded by dotted lines. The kNN distances were normalized by that of (a) SAP folds or (b) αβ class at k=100. The CATH fold class SP was not analysed due to its limited number among SAP folds which did not allow the calculation of nearest distance beyond k=10.

**Fig. S4: Additional novel folds, random coil content, and fold stability of SAP and DAP folds. a,** Additional representative SAP folds with similar SAP-like folds co-existed in the TED novel fold dataset. Structures were colored by the pLDDT scores assigned by ESMFold. Structures were sorted in descending order of the number of residues from the top left to the bottom right corner in each specific SAP/DAP-CATH fold class subpanel. Prefixes of the fold names were the COMPARE database IDs for the original APs. Suffixes of the fold names indicate the start and the end residue position of the AP folds in the original AP sequences. **b,** Random coil content and fold stability of SAP and DAP folds. DAP folds have statistically significantly higher random coil content ($p<0.001$) and higher conformational fluctuation of their ensemble backbones ($p<0.001$) than SAP folds. Two-tailed, independent Student's t-test was used for statistical testing.

**Fig. S5: Kingdom-level heatmaps showing the abundance of hit kingdoms of (a) SAP-like and (b) DAP-like folds in afdb50.** Columns represent the SAP and DAP folds origin kingdoms. Rows represent SAP-like and DAP-like folds hit kingdoms in afdb50. The color of each cell represents the count of origin-hit linkages normalized for over- or under-representation of origin kingdoms and the number of existing kingdoms in afdb50.

**Fig. S6: B-cell epitope propensity and surface feature comparisons among SAP, DAP, and SNAP folds. a,** Distributions of summed B-cell epitope scores normalized with summed epitope SASA of sized B-cell epitopes, which indicates the global B-cell epitope propensity of the folds. DAP folds showed an upshifted B-cell epitope propensity than SAP, SNAP and human proteome folds. **b,** Intercluster distances of SAP and food SNAP from human SNAP folds in the LDA-projected space of physicochemical profiles of sized B-cell epitopes assigned by DiscoTope and 3DZD. **c,** Top important physicochemical descriptors ranked by SHAP value that contributed to the class separation among SAP, food SNAP, and human SNAP folds in the LDA of Fig. S4d.

**Fig. S7: Projected 2-D maps of the orientation-specific differences in physicochemical properties of SAP and food SNAP from human SNAP folds.** The physicochemical properties were assigned on the entire fold surfaces independently by SURFMAP. Blue and red colors indicate the lower and higher physicochemical value compared to the human SNAP folds respectively. Sequence redundancy reduction to 90%, and re-sampling by bootstrapping (1,000 iterations) were applied to account for the effect of surface features redundancy and the population size differences among fold classes.

**Fig. S8: AllerX features important to the prediction of unseen APs. a,** SHAP value ranking for the features with the highest SHAP values in the prediction of each test set. IT is the internal test set. C24H/C25H and C24N/C25N are the subsets of seen APs with detectable sequence alignment hit (H) and unseen APs with no detectable sequence alignment hit (N) in COMPARE 2023, respectively. **b,** Distribution of summed bit score of sequence alignment >90% identity between human sourced HLA class II peptides (left) and the nonhuman sourced HLA class II peptides (right) to the polypeptide segments of NAPs and AP subsets. Regardless of the source, the summed bit scores were notably lower in the unseen AP subsets C24N and C25N compared to C23. **c,** SHAP value correlation with the feature values of esm_15 and progres_063 in the prediction of C24N and C25N. A higher SHAP value indicates the stronger push to the model to bias towards the positive class (i.e. allergenic) due to the feature value of a particular AP as a scatter dot. Both features presented a negative correlation to the SHAP value while not intercorrelated with each other as represented by the color of scatter dots.

**Fig. S9: Count of HLA class II peptide assigned by NetMHCIIpan, and the sequence characteristics of 4-, 5-, 6-mer from sequence alignment. a,** Counts of strong binder (SB) and weak binder (WB) HLA class II peptide assigned by NetMHCIIpan for the selected DR and DQ alleles. **b-f,** counts and sequence characteristics of exact matching 4-, 5-, and 6-mer derived from the hit HLA class II peptides. **b,** Number of the 6-mers from human (H) and nonhuman (NH) sources. The y-axis was normalized by the same method as Figs. 5a and b. **c,** Sequence space comparison of the 6-mers supervised for class separation among SAP, DAP, food SNAP, human SNAP, and human proteome folds by LDA on esm2 embeddings. **d,** Position-wise cos similarity of amino acid frequency from the UniProtKB background among the top 10% most abundant of the 6-mers. A lower cos similarity value indicates a more dissimilar amino acid frequency from the background. **e,** Sequence logos of the top 10% most abundant of the 5- and 6-mers in SAP, DAP, and food SNAP. **f,** Counts of the most abundant, exact matching 4-mers derived from the hit HLA class II peptides in food SNAP and human proteome folds. The counts were normalized by the same method as in Figs. 5a and b, and scaled by a factor of $10^{11}$ for readability.

**Fig. S10: Estimated binding likelihoods of the top 10% abundant HLA class II 4-mers with HLA-DR and HLA-DQ.** The binding likelihoods were expressed by the mean PSSM score of the simulated bound positions at P1-P6 positions in N➔C and the alternative C➔N orientations. Negative position numbers at the x-axis indicates the priming with an unbound N-terminal of the k-mer. For example, -1 means one amino acid residue in the k-mer N-terminal was unbound. Bracketing these scenarios of unbound k-mer N-terminals was for comprehensiveness of the analysis because the C-terminals of 5- and 6-mers when primed at P6 were also unbound, leaving a maximum of 2 C-terminal residues flanking beyond P9.

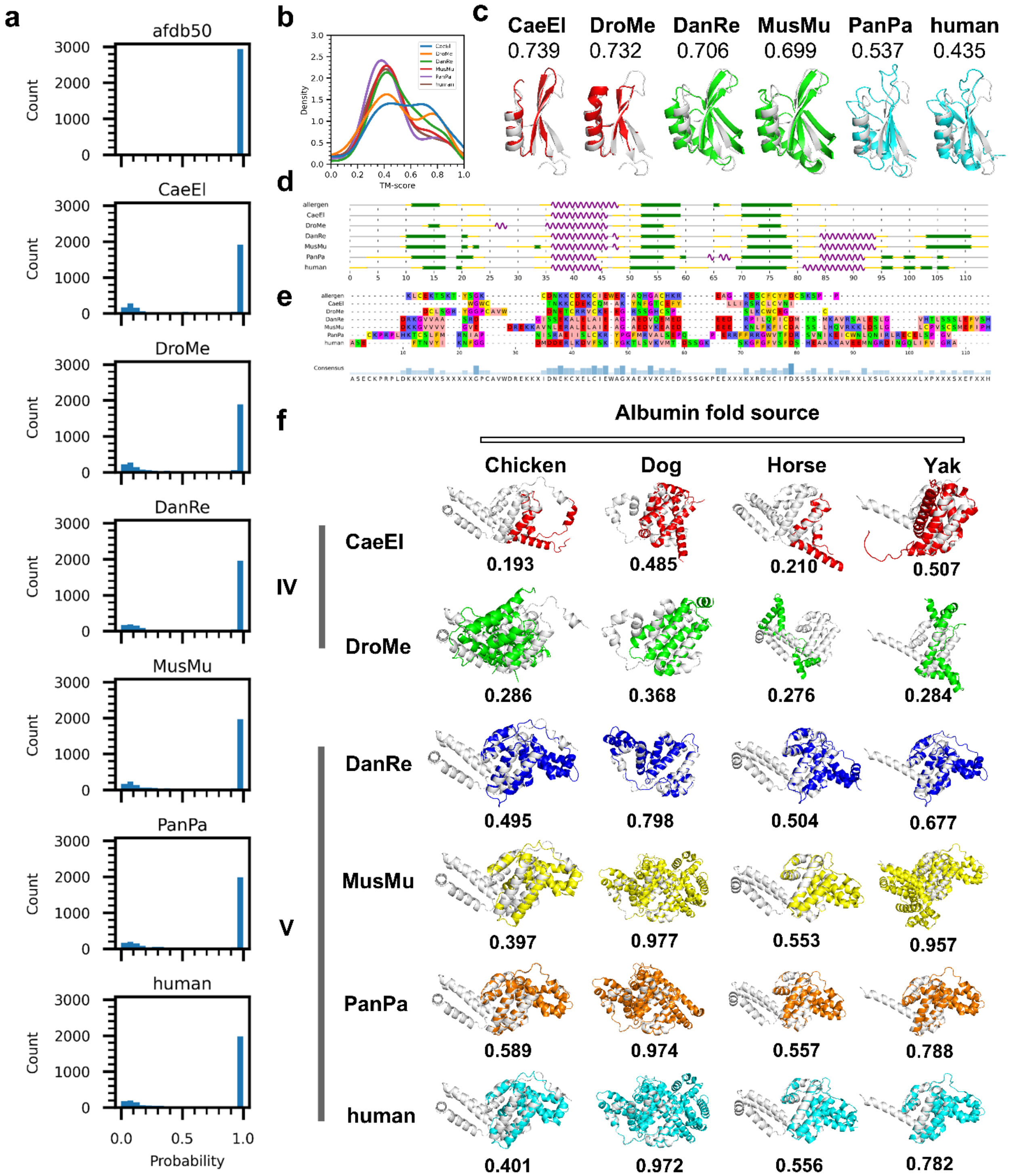
a
afdb50
CaeEl
DroMe
DanRe
MusMu
PanPa
human
Count
3000
2000
1000
0
0.0
0.5
1.0
Probability
b
Density
TM-score
c
CaeEl 0.739
DroMe 0.732
DanRe 0.706
MusMu 0.699
PanPa 0.537
human 0.435
d
e
Consensus
f
Albumin fold source
Chicken
Dog
Horse
Yak
IV
V
CaeEl 0.193 0.485 0.210 0.507
DroMe 0.286 0.368 0.276 0.284
DanRe 0.495 0.798 0.504 0.677
MusMu 0.397 0.977 0.553 0.957
PanPa 0.589 0.974 0.557 0.788
human 0.401 0.972 0.556 0.782

**Fig. S1**

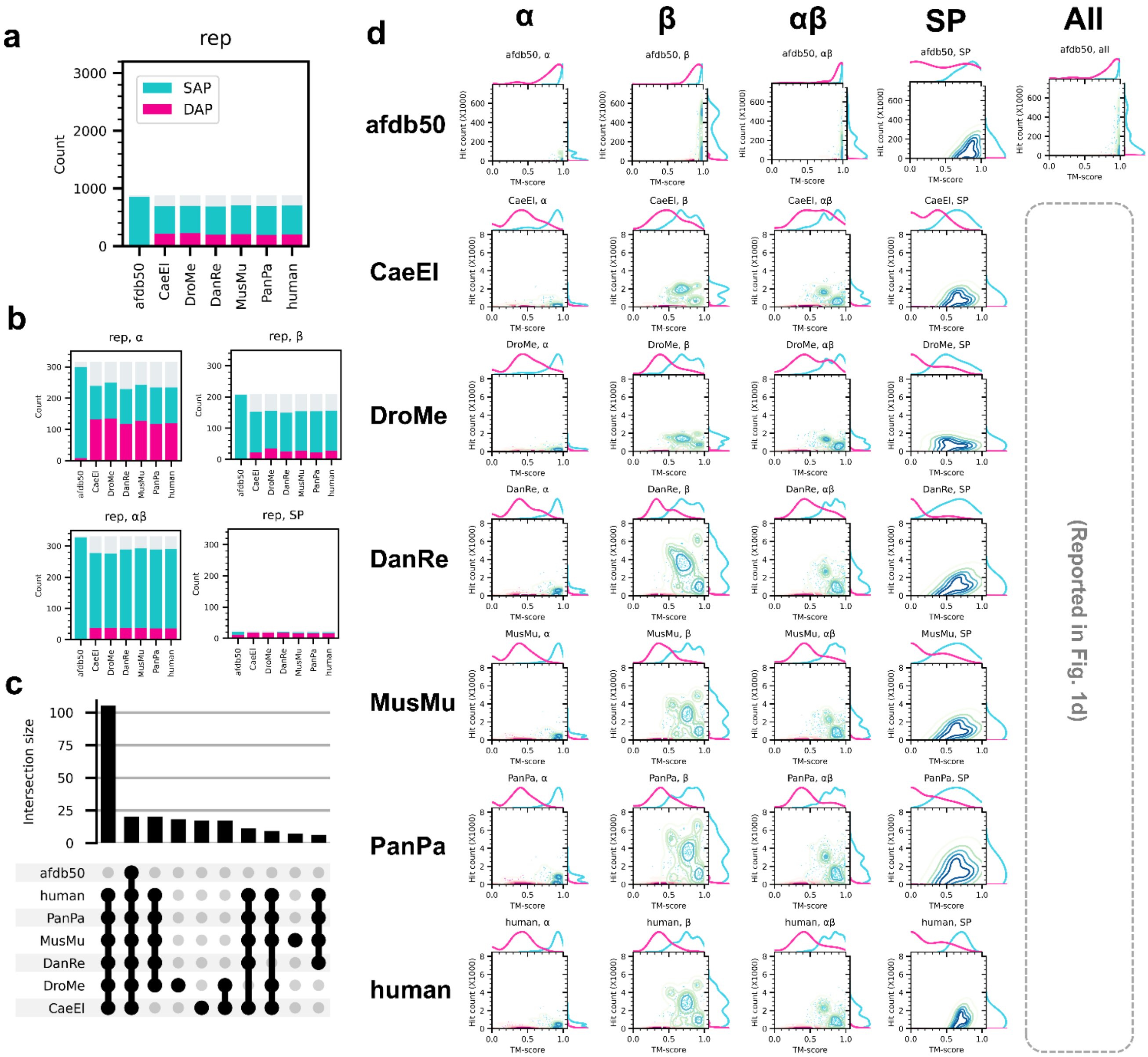

a
rep
SAP
DAP
Count
afdb50
CaeEl
DroMe
DanRe
MusMu
PanPa
human
b
rep, α
rep, β
rep, αβ
rep, SP
c
Intersection size
d
α
β
αβ
SP
All
TM-score
Hit count (X1000)
(Reported in Fig. 1d)


**Fig. S2**

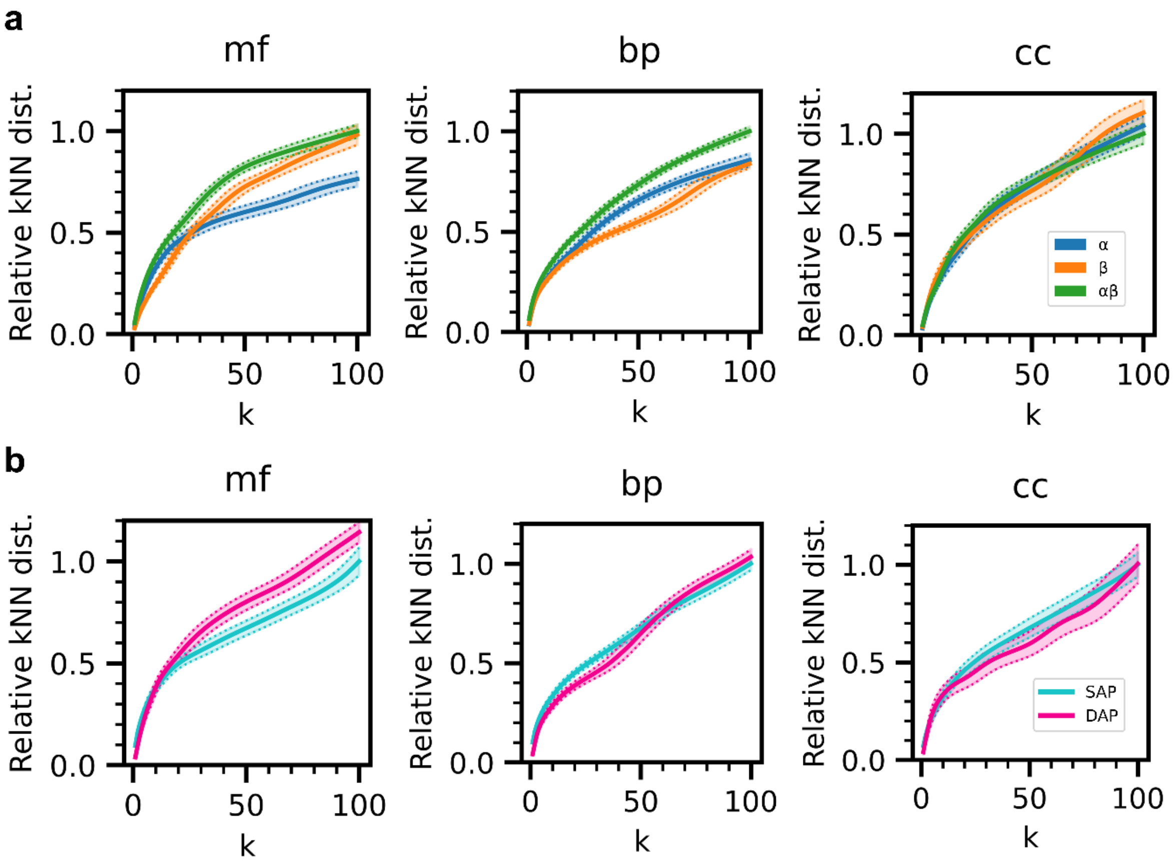


**Fig. S3**

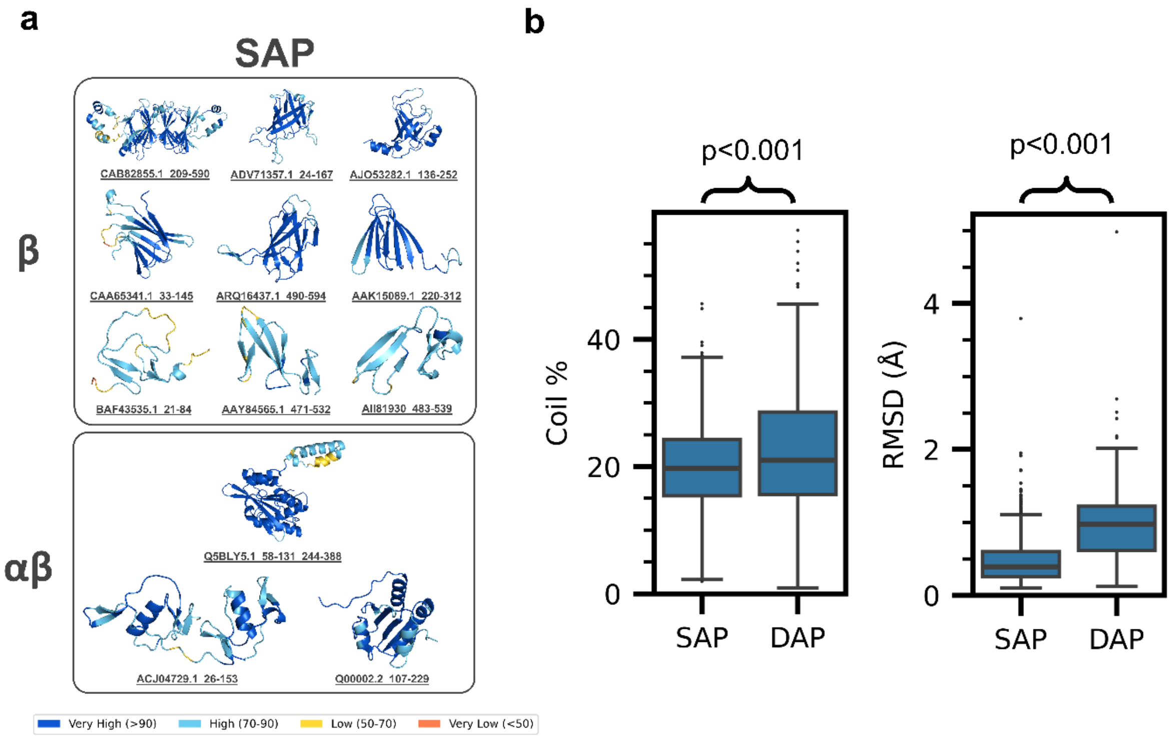
a
SAP
β
CAB82855.1_209-590
ADV71357.1_24-167
AJO53282.1_136-252
CAA65341.1_33-145
ARQ16437.1_490-594
AAK15089.1_220-312
BAF43535.1_21-84
AAY84565.1_471-532
AII81930_483-539
αβ
Q5BLY5.1_58-131_244-388
ACJ04729.1_26-153
Q00002.2_107-229
Very High (>90)
High (70-90)
Low (50-70)
Very Low (<50)
b
p<0.001
p<0.001
Coil %
RMSD (Å)
0
20
40
0
2
4
SAP
DAP
SAP
DAP

**Fig. S4**

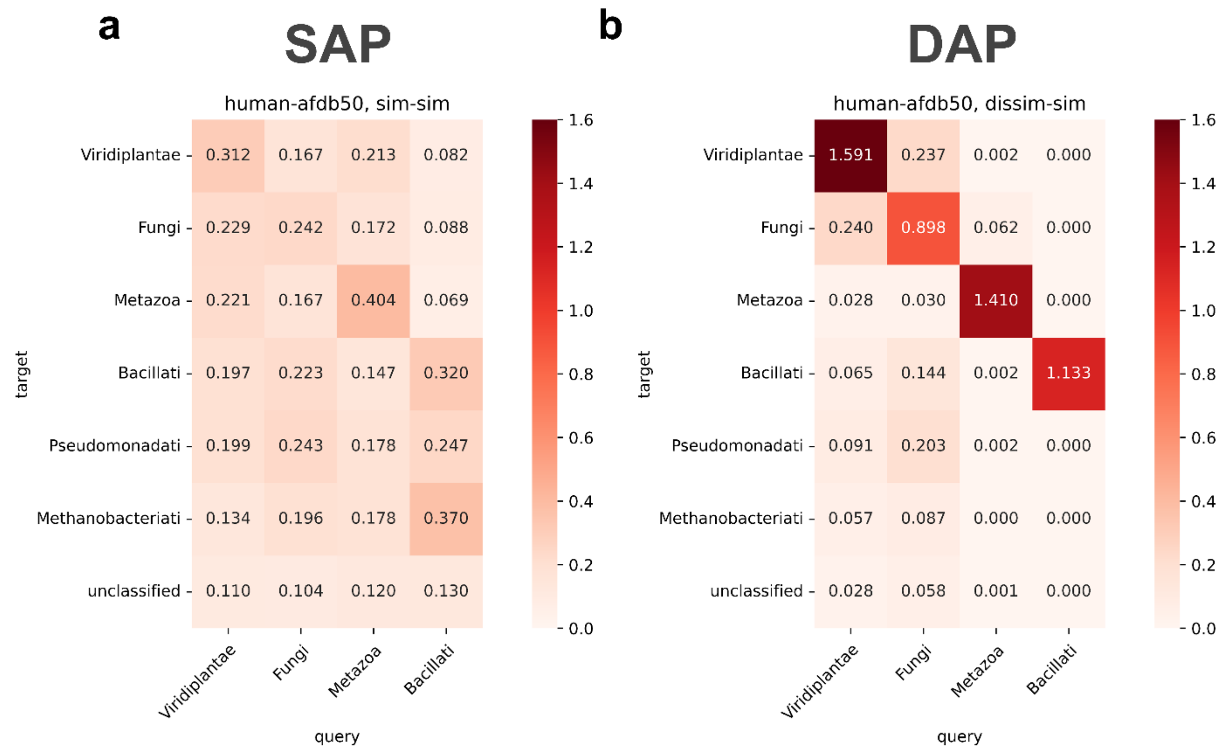
a
SAP
human-afdb50, sim-sim
Viridiplantae 0.312 0.167 0.213 0.082
Fungi 0.229 0.242 0.172 0.088
Metazoa 0.221 0.167 0.404 0.069
Bacillati 0.197 0.223 0.147 0.320
Pseudomonadati 0.199 0.243 0.178 0.247
Methanobacteriati 0.134 0.196 0.178 0.370
unclassified 0.110 0.104 0.120 0.130
target
Viridiplantae Fungi Metazoa Bacillati
query
1.6 1.4 1.2 1.0 0.8 0.6 0.4 0.2 0.0
b
DAP
human-afdb50, dissim-sim
Viridiplantae 1.591 0.237 0.002 0.000
Fungi 0.240 0.898 0.062 0.000
Metazoa 0.028 0.030 1.410 0.000
Bacillati 0.065 0.144 0.002 1.133
Pseudomonadati 0.091 0.203 0.002 0.000
Methanobacteriati 0.057 0.087 0.000 0.000
unclassified 0.028 0.058 0.001 0.000
target
Viridiplantae Fungi Metazoa Bacillati
query
1.6 1.4 1.2 1.0 0.8 0.6 0.4 0.2 0.0

**Fig. S5**

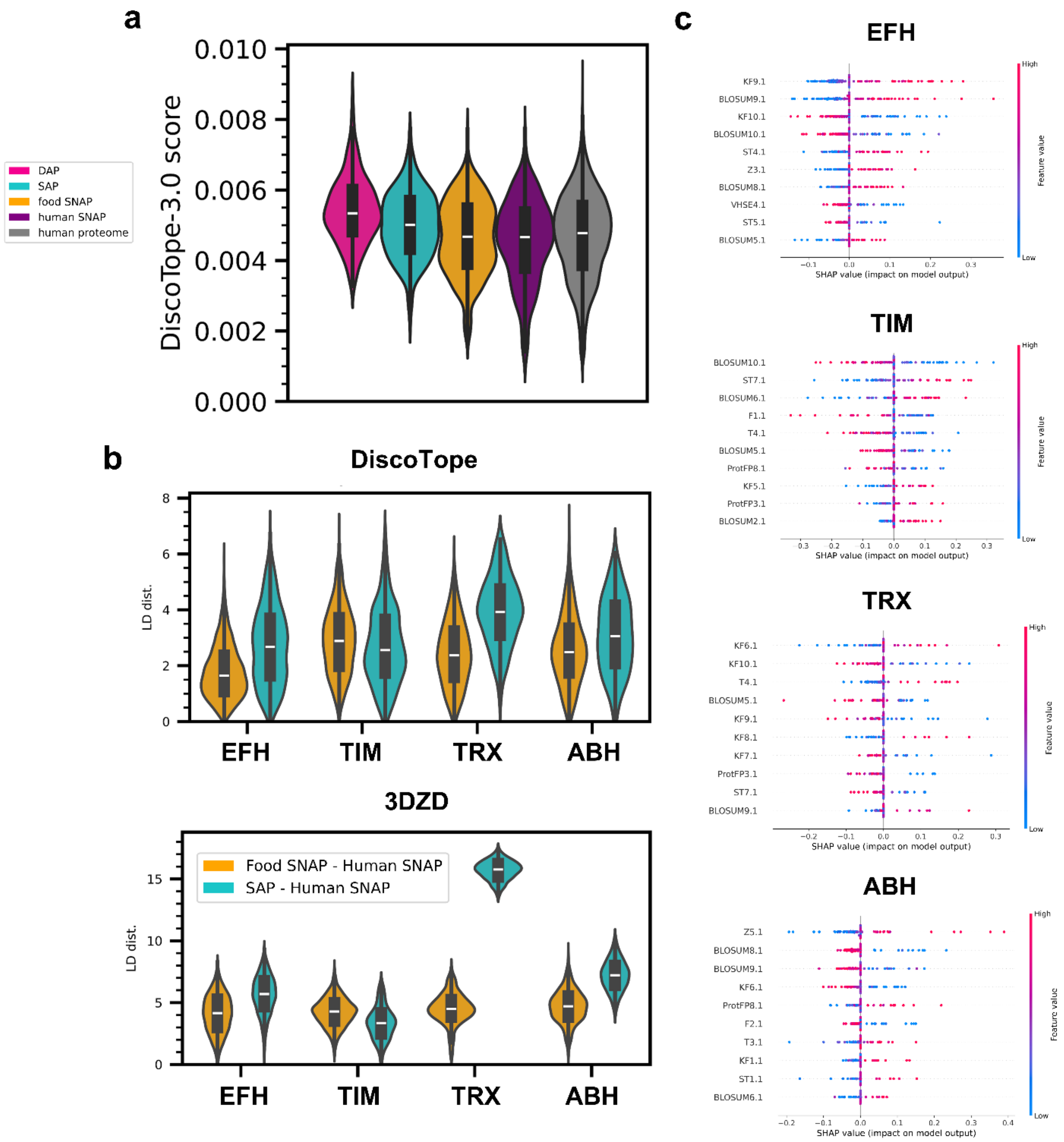
a
DAP
SAP
food SNAP
human SNAP
human proteome
DiscoTope-3.0 score
0.010
0.008
0.006
0.004
0.002
0.000
b
DiscoTope
LD dist.
EFH
TIM
TRX
ABH
3DZD
Food SNAP - Human SNAP
SAP - Human SNAP
c
EFH
TIM
TRX
ABH
SHAP value (impact on model output)
Feature value
High
Low

**Fig. S6**

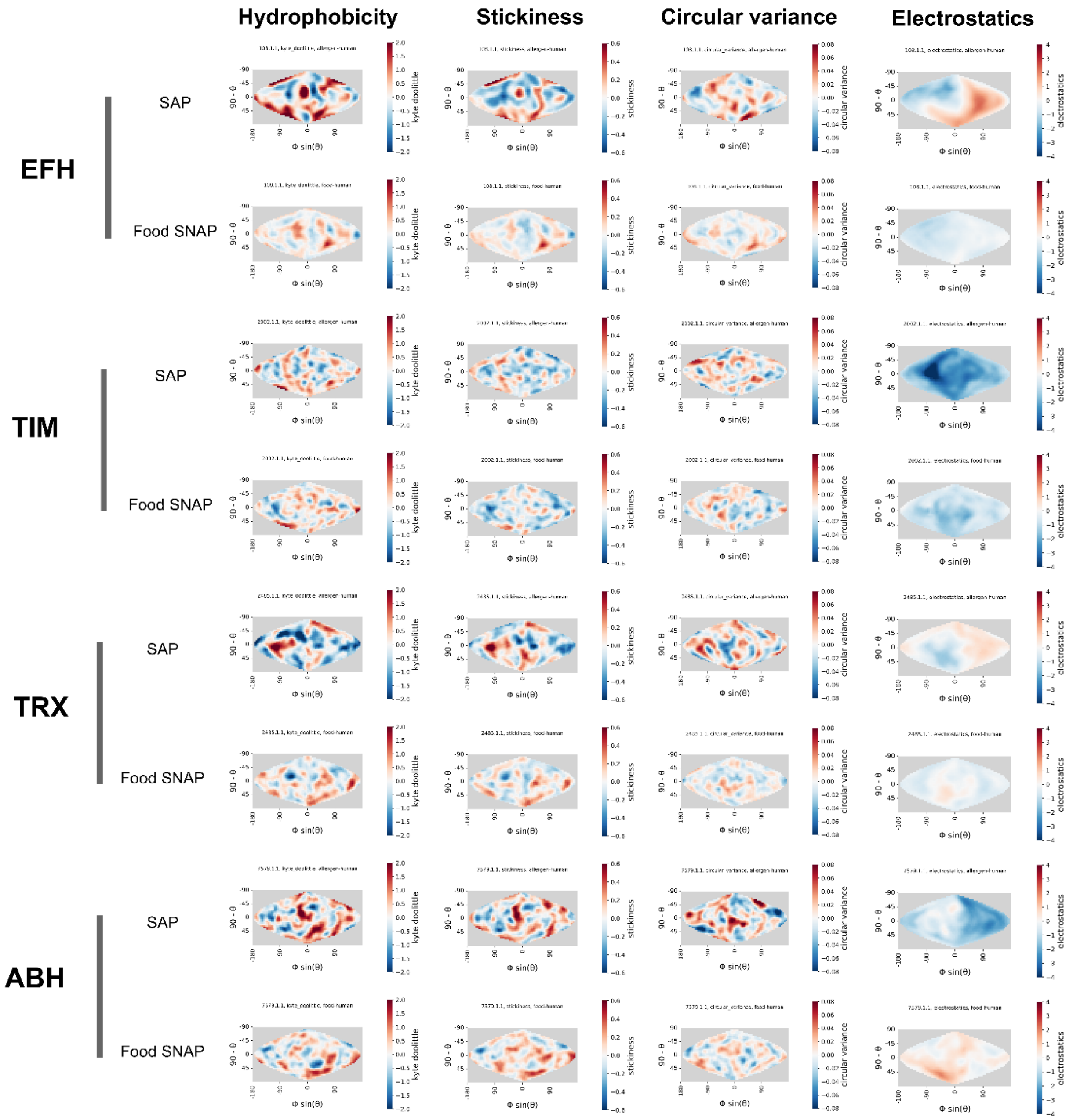
Hydrophobicity
Stickiness
Circular variance
Electrostatics
EFH
SAP
Food SNAP
TIM
SAP
Food SNAP
TRX
SAP
Food SNAP
ABH
SAP
Food SNAP

**Fig. S7**

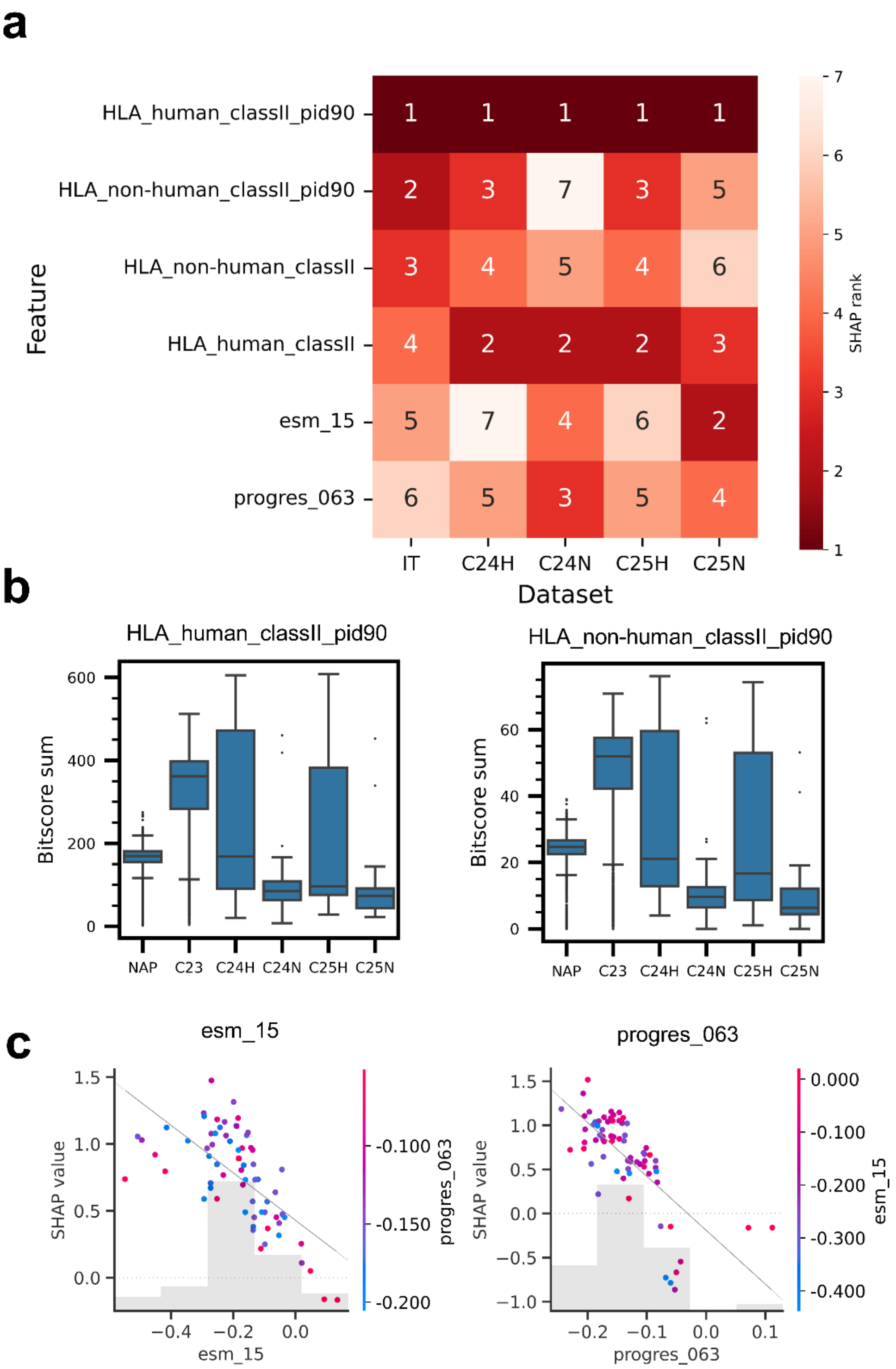
a
HLA_human_classII_pid90
HLA_non-human_classII_pid90
HLA_non-human_classII
HLA_human_classII
esm_15
progres_063
Feature
IT
C24H
C24N
C25H
C25N
Dataset
SHAP rank
b
HLA_human_classII_pid90
HLA_non-human_classII_pid90
Bitscore sum
NAP
C23
C24H
C24N
C25H
C25N
c
esm_15
progres_063
SHAP value

**Fig. S8**

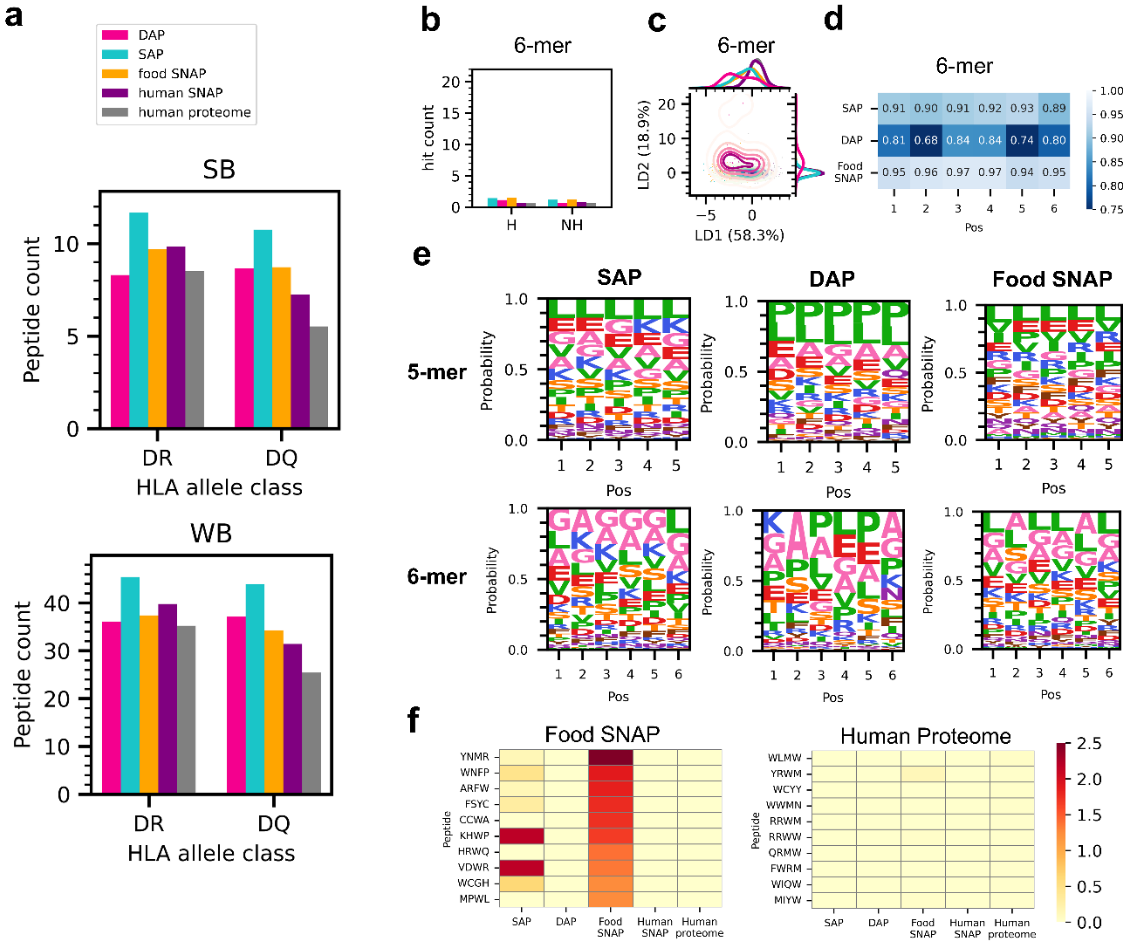
a
DAP
SAP
food SNAP
human SNAP
human proteome
SB
Peptide count
DR
DQ
HLA allele class
WB
b
6-mer
hit count
H
NH
c
6-mer
LD2 (18.9%)
LD1 (58.3%)
d
6-mer
SAP 0.91 0.90 0.91 0.92 0.93 0.89
DAP 0.81 0.68 0.84 0.84 0.74 0.80
Food SNAP 0.95 0.96 0.97 0.97 0.94 0.95
Pos
e
SAP
DAP
Food SNAP
5-mer
6-mer
Probability
Pos
f
Food SNAP
YNMR
WNFP
ARFW
FSYC
CCWA
KHWP
HRWQ
VDWR
WCGH
MPWL
Human Proteome
WLMW
YRWM
WCYY
WWMN
RRWM
RRWW
QRMW
FWRM
WIQW
MIYW
Peptide
SAP
DAP
Food SNAP
Human SNAP
Human proteome

**Fig. S9**

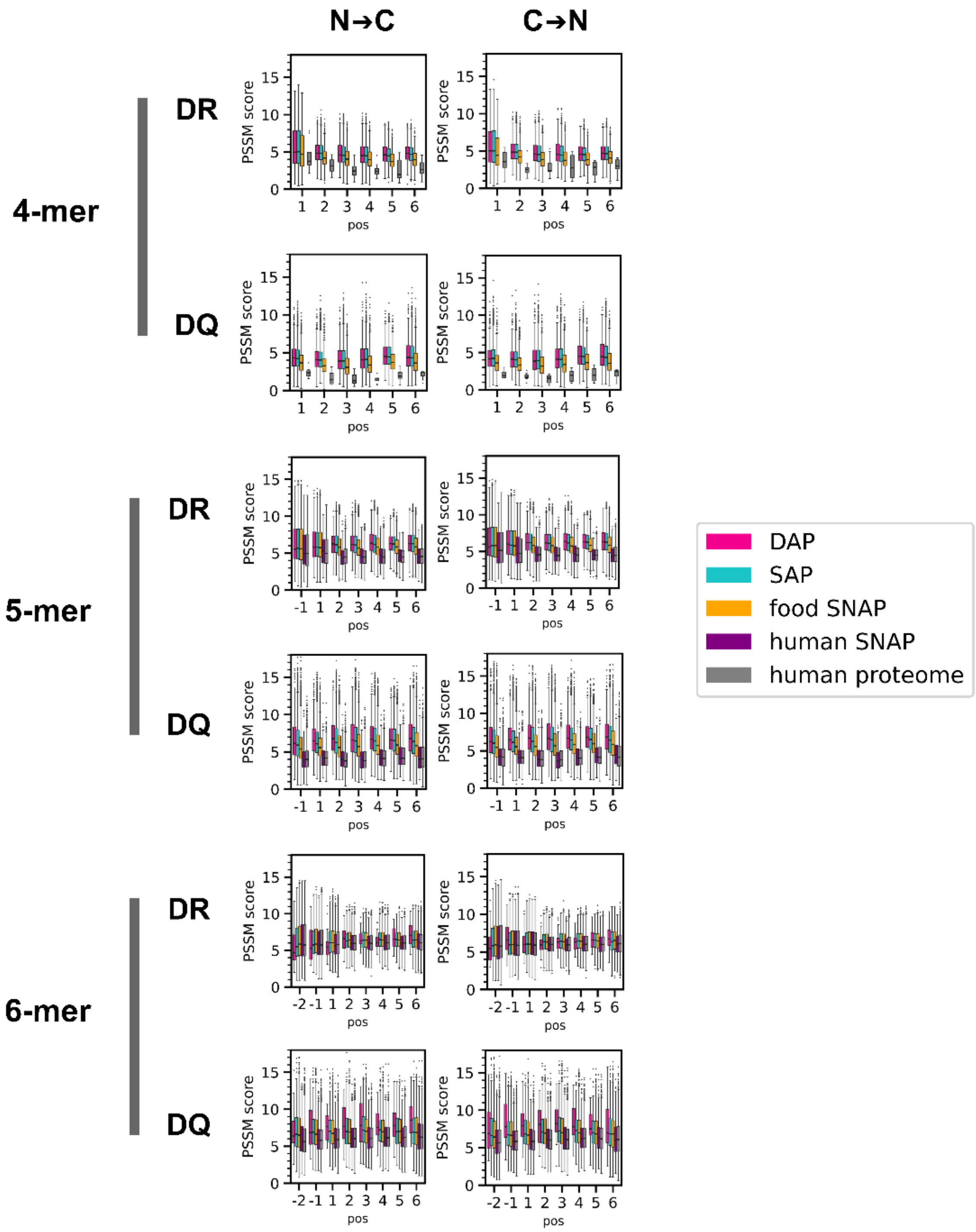
N→C
C→N
4-mer
5-mer
6-mer
DR
DQ
PSSM score
pos
DAP
SAP
food SNAP
human SNAP
human proteome

**Fig. S10**
